# Transcription, structure, and organoids translate time across the lifespan of humans and great apes

**DOI:** 10.1101/2022.10.28.513899

**Authors:** Christine J. Charvet, Kwadwo Ofori, Carmen Falcone, Brier A. Rigby-Dames

## Abstract

How the neural structures supporting human cognition develop and arose in evolution is an enduring question of interest. Yet, we still lack appropriate procedures to align ages across primates, and this lacuna has hindered progress in understanding the evolution of biological programs. We generated a dataset of unprecedented size consisting of 573 time points from abrupt and gradual changes in behavior, anatomy, and transcription across human and 8 non-human primate species. We included time points from diverse human populations to capture within-species variation in the generation of cross-species age alignments. We also extracted corresponding ages from organoids. The identification of corresponding ages across the lifespan of 8 primate species, including apes (e.g., orangutans, gorillas) and monkeys (i.e., marmosets, macaques) reveal that some biological pathways are extended in humans compared with some non-human primates. Particularly, the human lifespan is unusually extended relative to studied nonhuman primates demonstrating that very old age is a phase of life in humans that does not map to other studied primate species. More generally, our work prompts a reevaluation in the choice of a model system to understand aging given very old age in humans is a period of life with a clear counterpart in great apes.

**Significance Statement:** What is special about the duration of human development and aging has been an enduring source of interest. A significant hurdle in identifying which biological programs are unusually extended in humans is the lack of standardized approaches with which to align ages across species. We harnessed temporal variation in behavior, transcription, and anatomy to align ages across the lifespan of primates. These data reveal which biological programs are conserved, and which are modified. Harnessing time points across scales of study guides the choice of model systems to understand disease progression, and can be used to enhance care of great apes, many of which are critically endangered.

## Introduction

Humans differ in many aspects from other primates, though it is still unclear how biological programs have been modified in the human lineage. We know that developmental programs — such as synaptogenesis or brain growth — occur for an extended time in humans relative to primates (1–7). Biological changes and diseases that emerge in old age — such as brain atrophy or Alzheimer’s disease — have traditionally been elusive in non-human primates. This elusivity is either because these diseases are largely unique to humans or because they do not live sufficiently long lifespans for the disease to become manifest (8–10). A major hurdle in assessing which biological programs are conserved, and which have been modified in primates is the lack of a standardized approach with which to align ages across species (11). In the present study, we generated tools for cross-species age alignments, and we identified conserved and modified biological programs across primates.

The lack of great ape samples available for study has been a major hurdle in the study of human brain evolution. One solution to this problem has been to engineer organoids from human and great ape cells to identify conserved and modified developmental programs (12–14). Many but not all studies have converged on the finding that the pace of cell maturation is slower in humans than it is in non-human primates (15, 16). Whether cross-species variation in organoid maturational timelines relate to individuals remains an open question (17). Here, we integrate maturational rates of brain development across cellular and organismal scales to generate age alignments.

We captured a conspicuous number of time points (i.e., 573 time points) by aligning temporal changes in biological programs across multiple scales. The integration of time points from structural, behavioral, and transcriptional variation overcomes challenges of small sample sizes that are typical of primate studies. We used these data to identify corresponding ages across the lifespan of human and non-human primate species (18, 19). In doing so, we implemented machine learning models to generate corresponding ages (20, 21). We also included time points from diverse human populations and great apes living in different conditions (e.g., captive versus wild) to capture within species variation in extrapolated time points (22, 23). Comparative analyses of survival rates across diverse human populations show that the human lifespan is unusually extended compared with studied great apes.

## Results

### The Dataset to Translate Ages Across the Lifespan

We considered 573 time points to align ages across 9 primate species (e.g., humans, orangutans, gorillas, rhesus macaques, gibbons, and marmosets; Fig. 1–3; Table S1). Some of these are structural, behavioral, and transcriptional (Figs. 1–7; Figs. S1–S7), but also include life history. Time points were obtained across pre- and postnatal periods. They include the age at which prenatal reflexes emerge, and when neurons switch from a proliferative to a post-proliferative state. Postnatal time points were collected from abrupt and gradual changes in brain, body, and behavior (Figs. 1–4). Most of the data are from individuals, but approximately 2% of these time points are from the organoids (n=11; Figs. 4 and S5). We collected time points (e.g., age of menarche) from several human populations and from nonhuman primates living in different environments (i.e., wild or captive) to capture within species variation (e.g., Figs. 5A-D and S1). Most of the time points are from captive individuals (Fig. S1), and we considered sex differences (Fig. 5 E, F and Table S1).

**Fig. 1.**
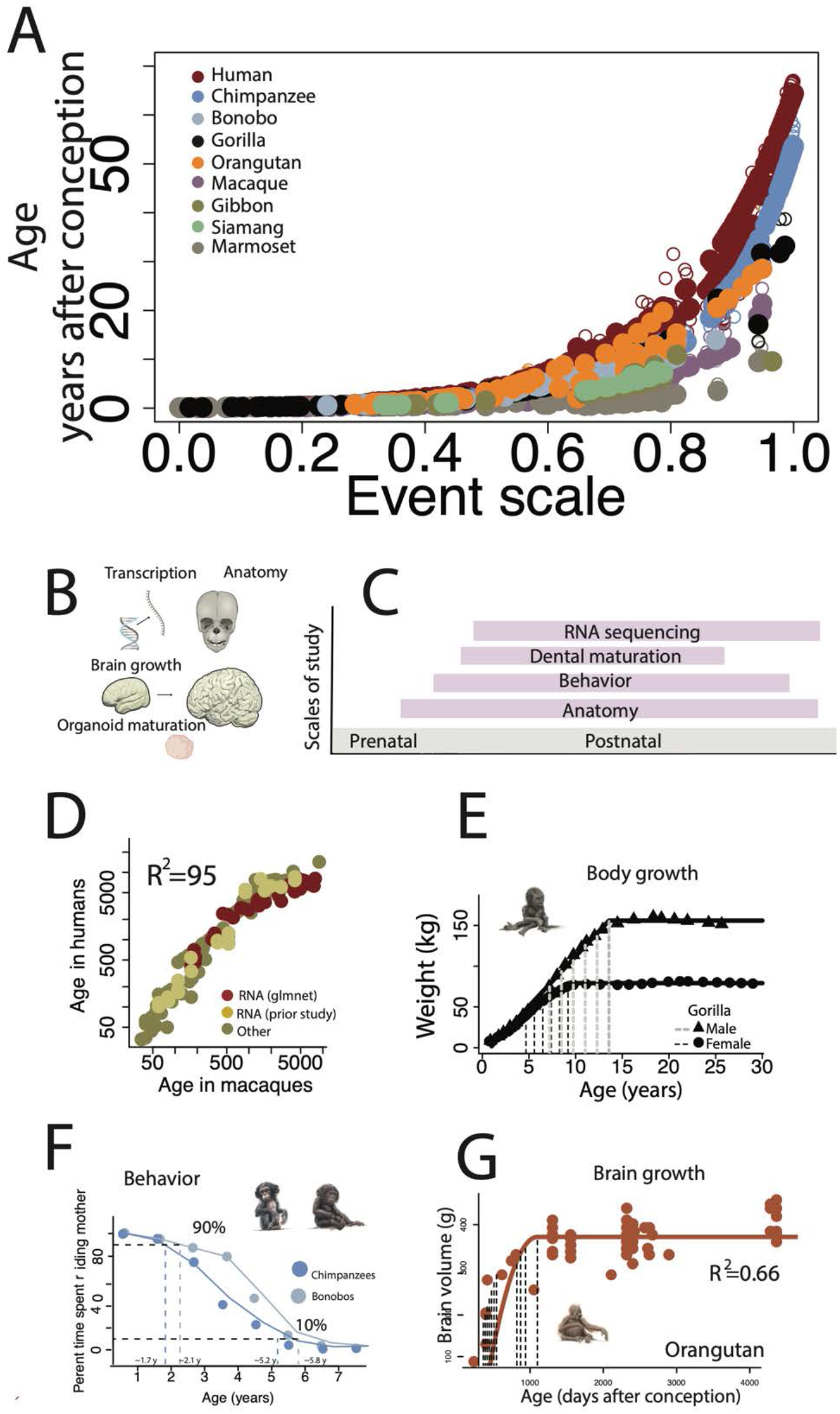
(A) We developed a model to find corresponding ages across species. Predicted (closed circles) and time points (open circles) are regressed against an ordering of time points (the event scale). (B) We used several metrics spanning a wide age range with different metrics represented on a life timeline. (D-G) We illustrate a few examples. (D) We captured time points from machine learning models from transcriptional variation. (E) We also extracted time points from non-linear regressions (E) applied to body growth as shown for a gorilla. We captured when the body reaches a percentage of adult volume (e.g., 100%, and 90%; vertical bars). We also fit smooth splines (F) through time spent riding mothers in chimpanzees and bonobos and we quantified when the values reach a percentage of time spent riding. (G) We also considered brain growth and collected when the brain reaches percentages of adult volume (vertical bars) as shown in orangutans.

**Fig. 2.**
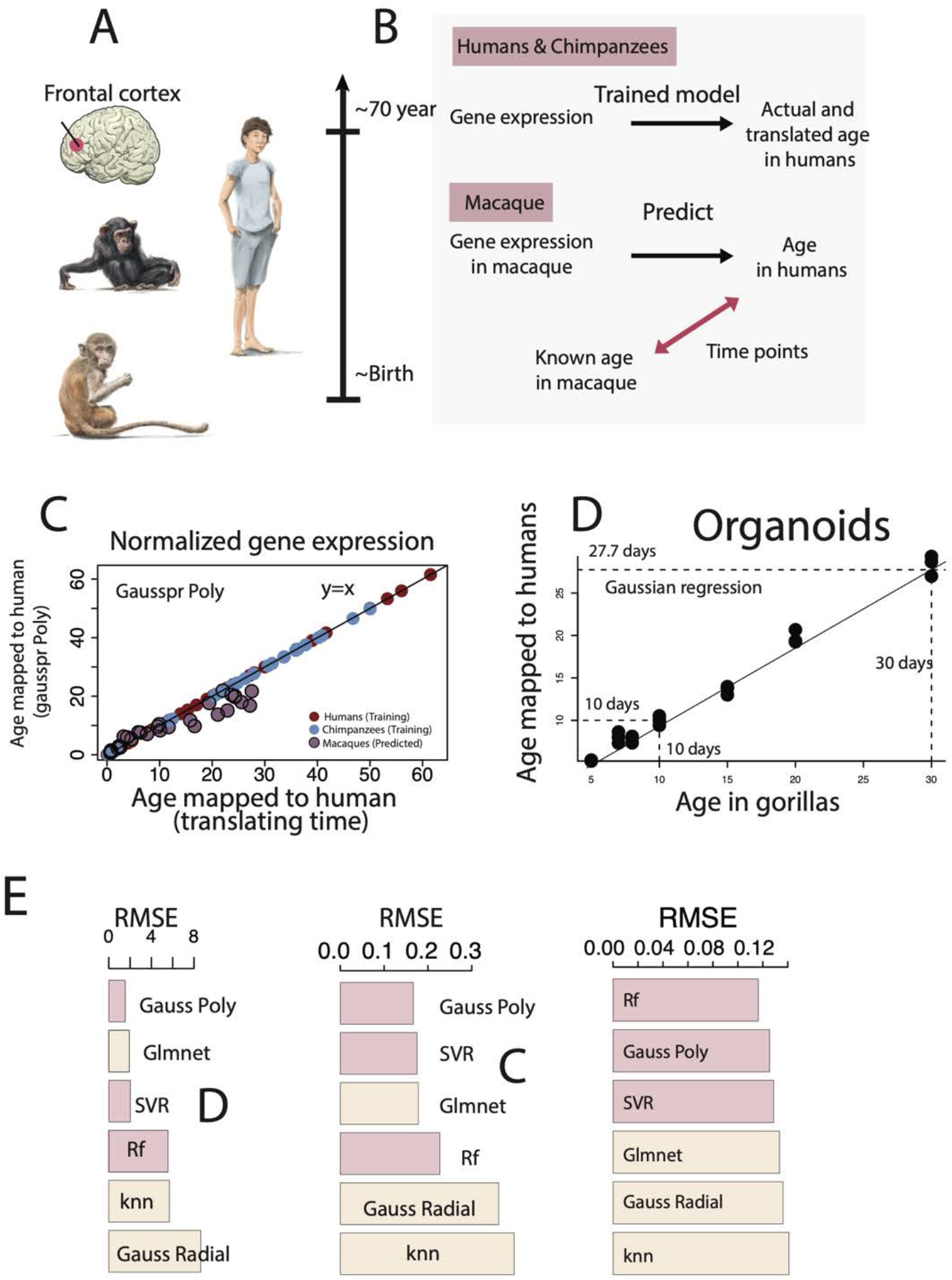
(A) We generated cross-species age alignments from frontal cortex normalized gene expression in humans, chimpanzees, macaques and organoids. (B) We trained the model to predict age in a set of species. We selected the model yielding the highest age prediction accuracy (root mean square error: RMSE). We then used the selected trained model to predict age based on normalized gene expression from individuals of a different species. Because the model had been trained to predict age in humans, the model predicted ages translated to humans. We used known ages of nonhuman primates and those translated to humans as time points in the model. We include examples of age alignments from tissue (C) and from brain organoids (D). (E) We tested the gaussian process regression (Gauss poly), gaussian radial regression (Gauss radial), random forest (rf), lasso, and elastic-net regularized generalized linear models (glmnet R package), support vector regression (SVR), and k-nearest neighbor (knn).

**Fig. 3.**
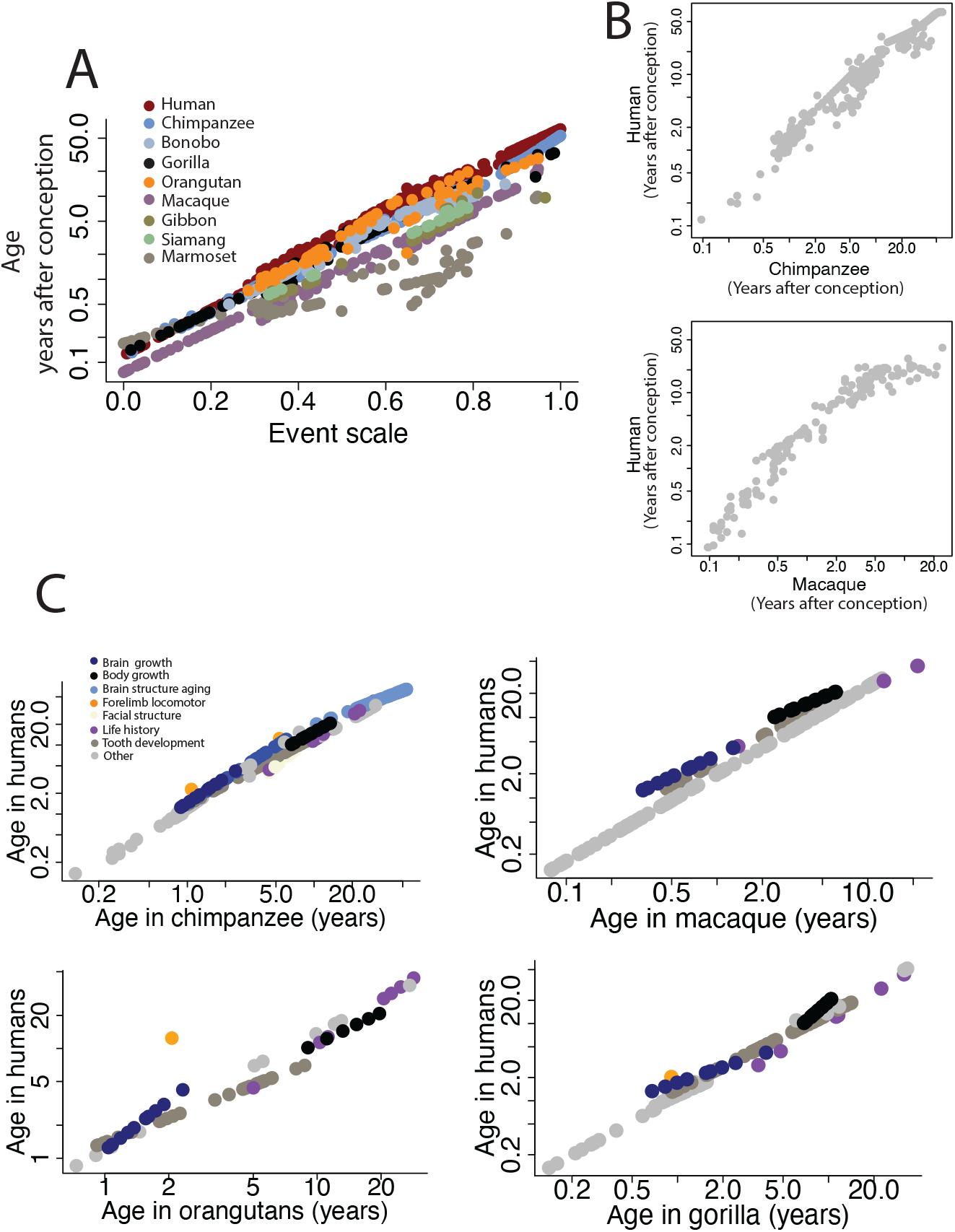
(A) Predicted time points expressed in years after conception are log-transformed and plotted against an event scale (an ordering of time points) for different species in the study. These analyses show that corresponding ages between humans and great apes are similar early in development but that corresponding ages gradually diverge with age across these taxa. Macaques and marmoset proceed at a pace of development and aging that differs from great apes and humans. Time points in macaques occur earlier than they do in great apes and humans. (B) We also include observed time points for two-species comparisons. (C) Output of the model shows that some translated time points deviate relative to others.

**Fig. 4.**
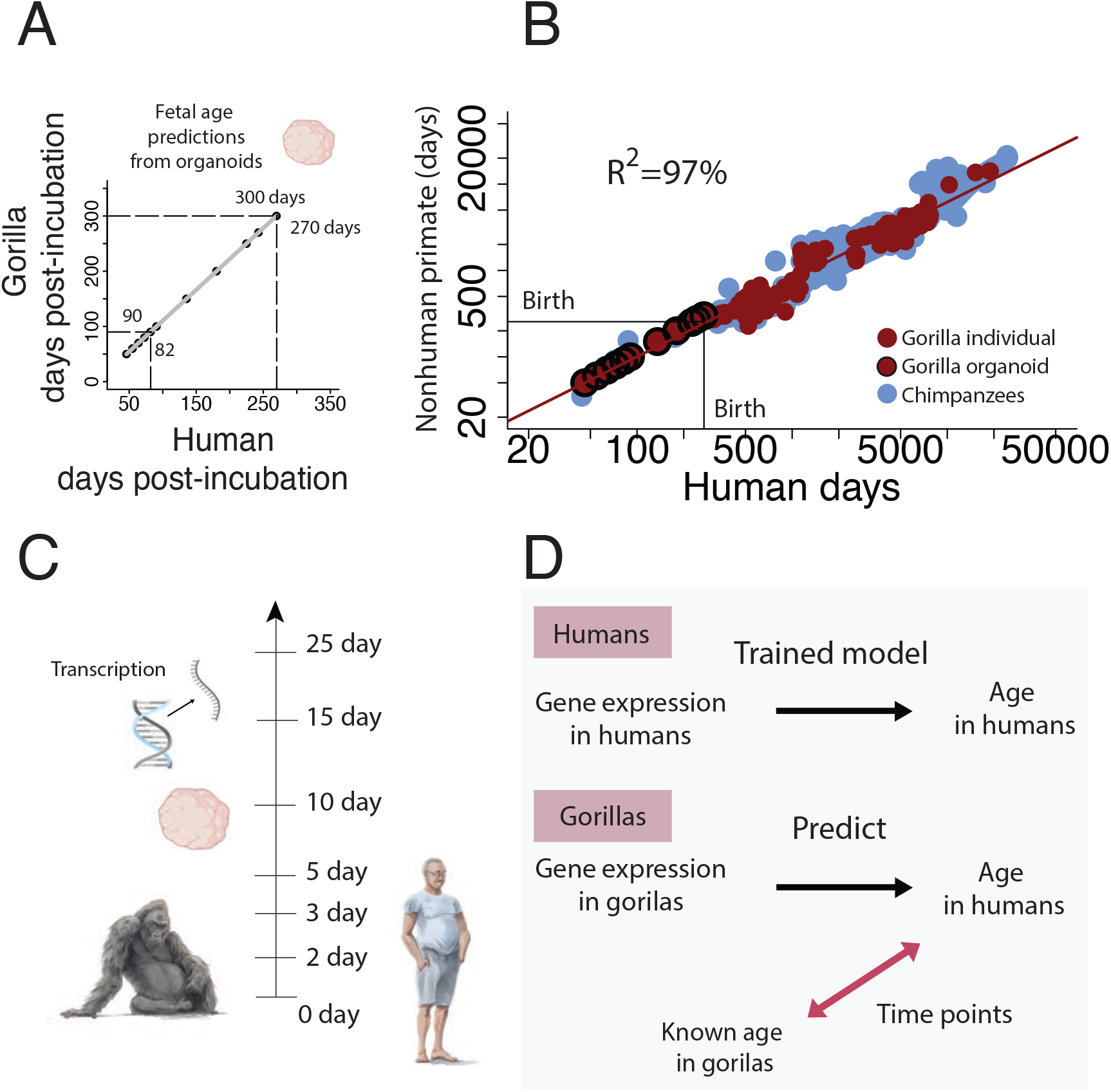
(A) We identified corresponding ages between human and gorilla organoids. (B) Cross-species age alignments from organoids were similar to those extracted from individuals. (C) We used organoid normalized gene expression extracted from 0 to 25 days after incubation onset. We used 7 different maturational states from both humans and gorillas to train machine learning models to align ages across the two species. (D) We first trained models to predict age in humans from human organoid gene expression. We then used this trained model, imported normalized gene expression collected from gorillas, and predicted age. The consequence of importing data from normalized gene expression in gorillas in these models, which had been trained in humans, is that the model outputs human age (i.e., age translated from gorillas to humans). We used the age of gorilla organoids and the age of human organoids as a basis with which to translate ages across species. (B) Time points collected from organoids align with time points collected from individuals.

**Fig. 5.**
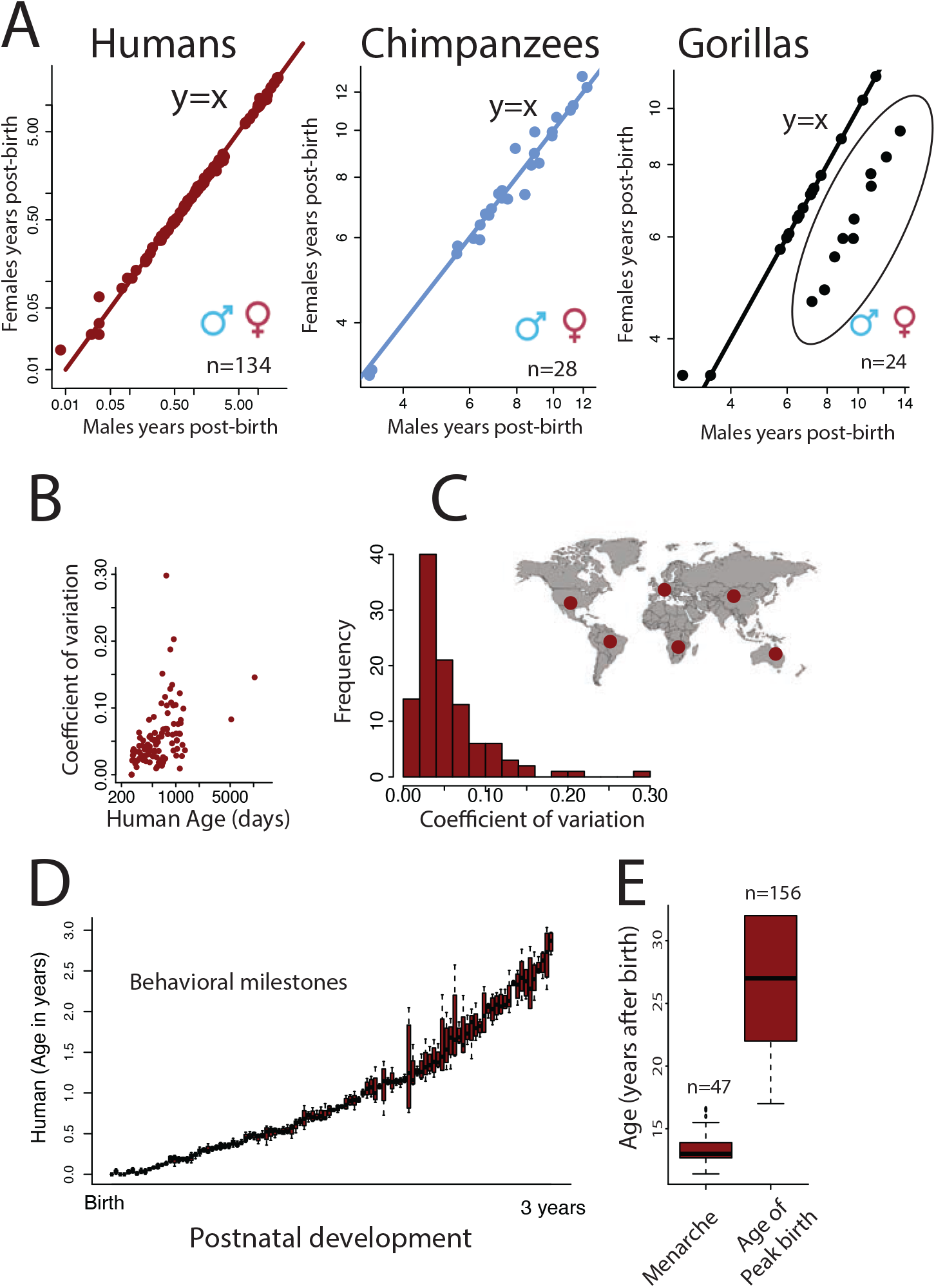
We considered individual variation (A-E) in collected time points in translated ages, including (A) sex differences. The pace of development is similar across males and females in humans, chimpanzees, and gorillas with most time points close to y=x. A few time points deviate from others as is evident in gorillas where body growth is protracted in males versus females (circles encapsulated by a sphere). Overall, (B) the coefficient of variation is similar across ages in humans though the standard deviation increases with age. This is evident when comparing locomotor milestones (D) across the first years of life and the distributions in the age of peak birth versus menarche in humans.

**Fig. 6.**
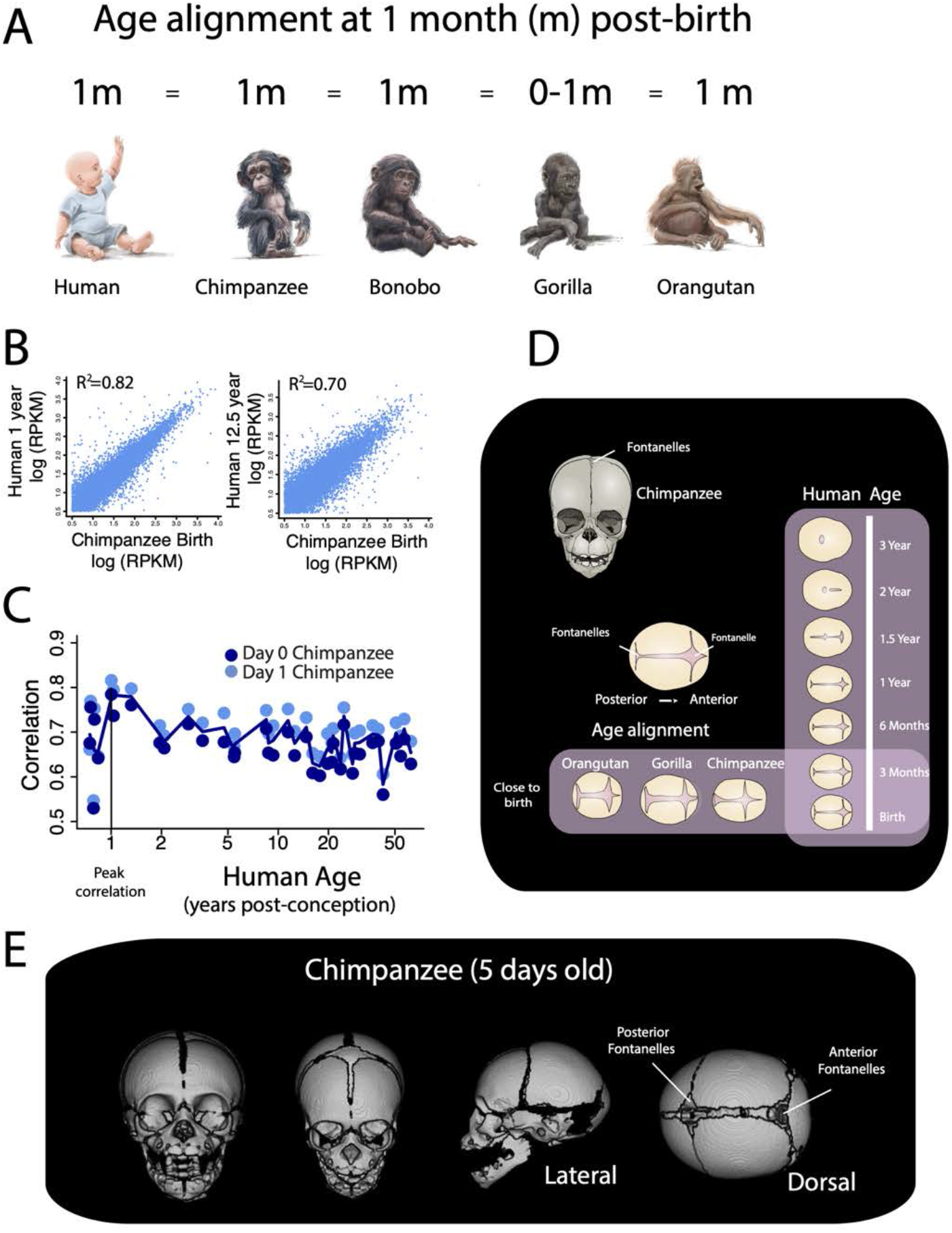
(A) According to our model, humans and great apes are similar in level of maturity around birth. A newborn roughly equates to a 1-month-old chimpanzee and a newborn orangutan. We also considered normalized gene expression (B-C) and fontanelles (D) as an additional basis to align ages around birth. (C) We correlated the log-transformed RPKM values of the frontal cortex of chimpanzees around birth (day 0 and 1 day after birth) with those of humans at different developmental ages. (C-D) The correlation coefficients from tested associations between the log-transformed RPKM values between the frontal cortical areas in chimpanzees near birth (at day 0 and day1) and those humans from different ages show that the frontal cortex of chimpanzees resembles humans that are slightly older than at birth. More precisely, newborn chimpanzees mostly align with humans at 1 year of age. (D) We next evaluated great ape and human fontanelles. We drew fontanelles from the dorsal view of the skull CT scans. The posterior fontanelle recedes followed by the anterior fontanelles with age in humans. The anterior and posterior fontanelles of orangutan, gorilla, and chimpanzee individuals close to birth resemble the fontanelles of newborn humans, which suggests great apes most closely resemble humans around birth. (E) CT scans used in are from the Digital Morphology Museum, Kyoto University Primate Research Institute (KUPRI). Anterior and posterior fontanelles are evident in this 5-day old chimpanzee.

**Fig. 7.**
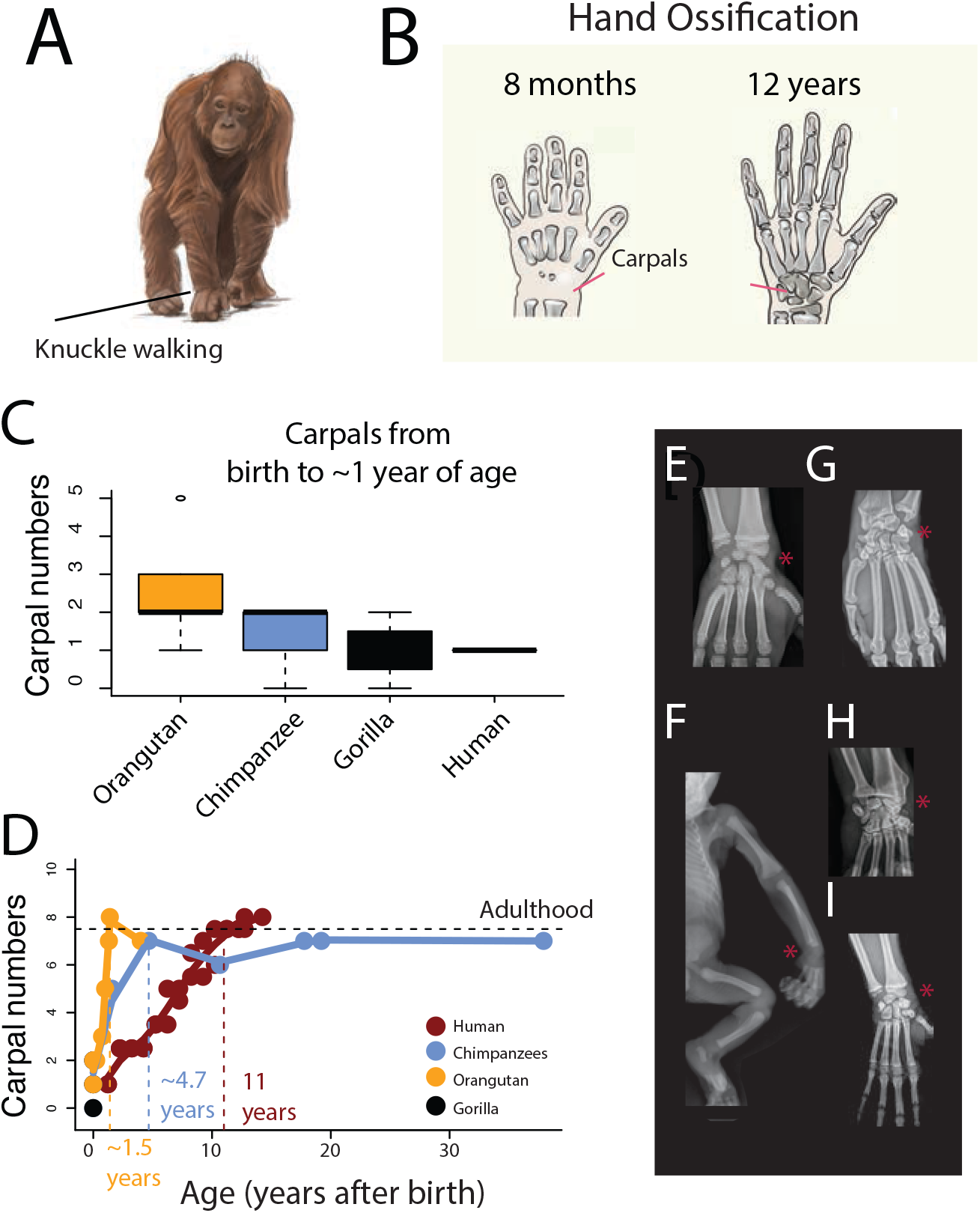
(A) Great ape knuckle-walk (e.g., orangutans; A) but not humans. We focused on carpal maturation (B) because carpals are substrates for knuckle walking. (C) We quantified ossified carpal numbers at birth across different species and over the course of postnatal maturation (D) from CT scans as well as radiographs from different great apes (E-I). Red asterisks highlight carpal bone ossification. (C) Within the first year of life, some great apes have a higher number of ossified carpal bones than humans, but these differences are not statistically significant. Species differences become increasingly evident postnatally. (D) Ossified carpal bone number increases at a much faster pace in great apes (i.e., orangutans, chimpanzees) than in humans. Adult carpal numbers are reached at approximately 1.5 years of age in orangutans, 4.7 years of age in chimpanzees, but 11 years of age in humans. We include some examples of radiographs from gorillas (E, I), including a newborn gorilla (F), and chimpanzees (G-H; 33).

### Machine Learning Models Generate Age Alignments from Brain Transcription

We used gene expression (Figs. S1–S7) and structural metrics (Figs. S8–S10) to test 6 machine learning models (i.e., lasso and elastic-net regularized generalized linear models, support vector regression, k-nearest neighbor, random forest, gaussian process regression) to predict ages across species. We used three datasets, two of which consisted of normalized gene expression extracted from the frontal cortex of individuals varying in age (Figs. 1–4, and Tables S1), and one of which consists of normalized gene expression from human and gorilla organoids also varying in age (n=42; Figs. 4 and S5). These data consist of relatively few individuals with many sampled genes (10,000~13,000). We selected 6 models that have made successful predictions with relatively small samples (20, 24–26; Figs. 1–3 and S4–S7). We randomly partitioned the data into a training set (~70% of the data) and a testing set (~30% of the data). We used a measure of the difference between predicted and observed ages, called the root mean square error (RMSE), to assess age prediction accuracy (Figs. S4–S7). We applied a similar approach to the generation of cross-species age alignments from diffusion metrics and growth trajectories (Figs. S8–S10).

For cross-species age alignments generated from transcription in tissues, the training set was drawn from humans and chimpanzees (Figs. 1–4 and S2–S4). We first translated age in chimpanzees to human age according to past work (19) and we trained these models to predict chronological age within species. Overall, RMSE values were similar across models (Fig. 2E), but the glmnet, support vector regression, random forest, and gaussian regression produced the lowest RMSE scores across datasets (Figs. 2E and S5–S7). We trained each model to predict age in humans. We then imported normalized gene expression from species other than humans to predict age. Because the model had been trained to predict age in humans, the model translates the age of imported normalized gene expression from non-human primates to human age. We used the ages of non-human primates and those translated to humans in the model.

We compared translated ages from machine learning models with past work (19, 27, 28). We log-transformed time points with age expressed in days after conception, and we fit a linear model to predict age in human from the age of macaque time points and their square (R2=0.99; F=979; p<0.01; Figs. 1, 3 and S4; Fig. 1D), but the inclusion of RNA from our machine learning model as a factor does account for a significant percentage of the variance (estimate=0.14 t=5.078, p1.03e-06). There is strong overlap in extrapolated ages across methods, which supports the notion that they are valid to generate cross-species age alignments.

### Machine Learning Models Generate Age Alignments from Organoids

We used machine learning models to find corresponding ages across human and gorilla organoids (Figs. 4A, B and S5) collected from 0 to 25 days post-incubation (Fig. 4C). We first trained the model to predict age in humans from gene expression. We then imported normalized gene expression in this trained model to predict age from gorilla organoids (Fig. 4D). Since the model had been trained to predict age in humans, inputting normalized gene expression from gorilla organoids predicts age in humans. We used the age of gorilla organoids and those translated to humans as corresponding time points in the model.

We compared cross-species age alignments from organoids with other metrics to determine how the use of organoids may apply to translating ages across species (Fig. 4B). We fit a linear model to log-transformed time points from organoids (n=11) and individuals (n=147) in humans and gorillas (with gorillas as the predictor variable) The model accounts for a significantly high percentage of the variance (y=1.08x-0.13; R2=97%). The addition of tissue type (in vivo versus in vitro) accounts for a significant percentage of the variance (estimate=-0.11; t=-2.9; p=0.004). Therefore, there are some differences in age alignments between organoids and individuals.

### A Model to Translate Ages Across the Lifespan

We used data from structural, transcriptional, and behavioral variation and we fit a general linear model to equate log-transformed corresponding ages across species. We first imputed the data because time points are not collected across all species. We ordered time points to generate an event scale. The event scale was generated by subtracting each time point (averaged across species) by the minimum age and dividing that time point by the difference between the maximum and minimum ages averaged across species. The ordering of time points ranged from 0 to 1 with early time points assigned a low score and later time points assigned a score close to 1. We included predictors and factors in the model. Those include the event scale, species, and event type. We use event type to test for heterochrony (i.e., modifications in the relative timing of biological processes) below. We tested the following factors:

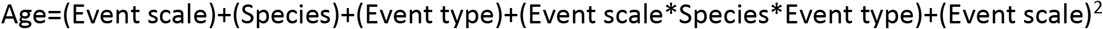

Age is log-transformed days after conception. This model accounts for a significantly large percentage of the variance (98.8%; F=978.7; p<2.2e-16; Figs. 1 and 3) and captures the pace of development and aging across species. Early in development, corresponding ages are roughly similar across humans and great apes, but corresponding time points gradually diverge with age (Fig. 3, 8). Humans and great apes are roughly similar in age in their first year of life, but a human in their mid-30s equates to a chimpanzee in their mid-twenties and a gorilla in their early 20s (Figs. 1, 3, and 8). The pace of aging is extended in humans compared with non-human primate species.

**Fig. 8.**
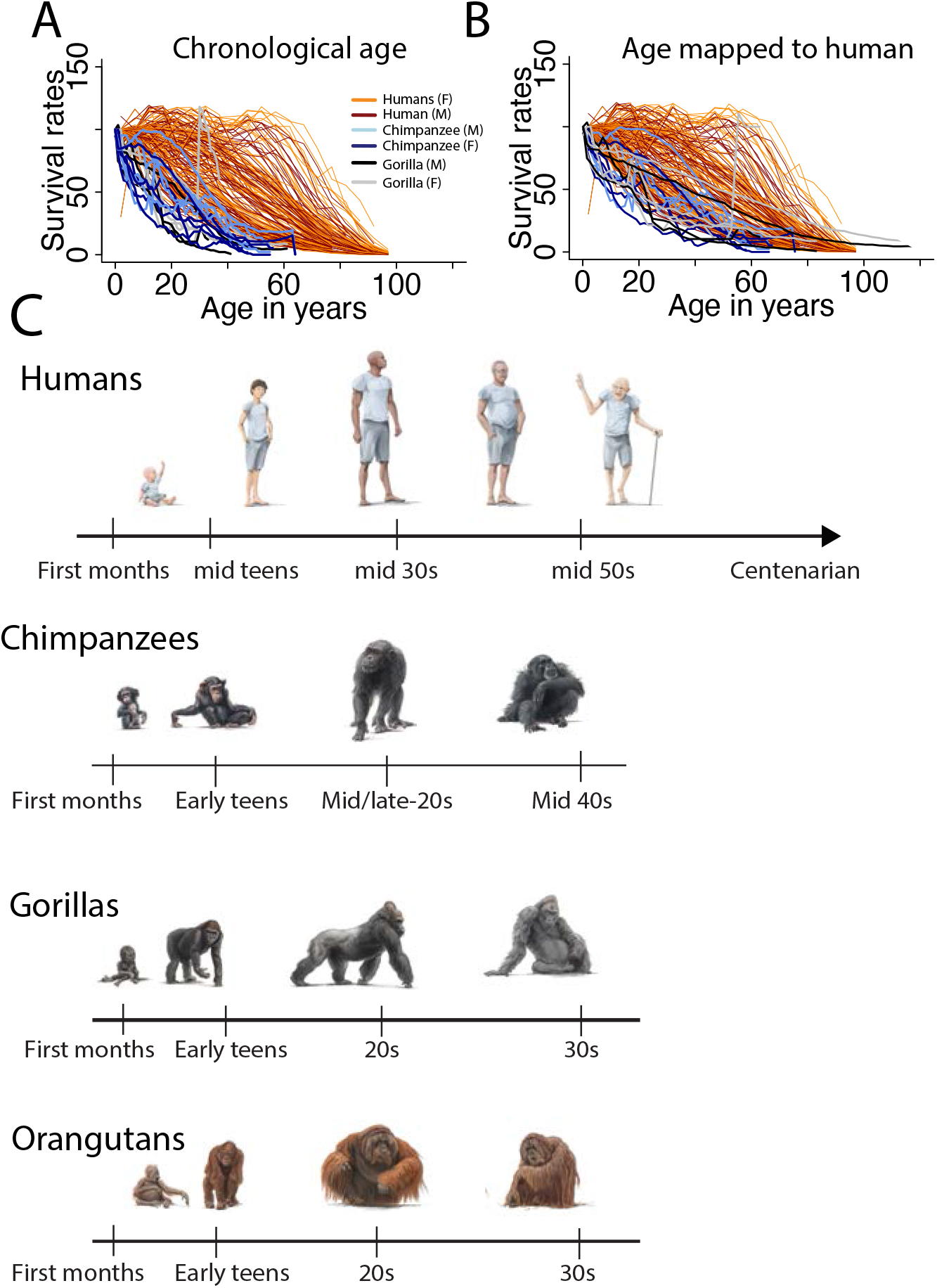
We captured survival rates across diverse human populations as well as humans and chimpanzees. Separate survival rates are generated for males and females (A) Survival rates are plotted against chronological age in humans and gorillas, which show that humans live overall longer than chimpanzees and gorillas in terms of absolute ages yet there are species differences in the pace of development and aging that should be controlled for. (B) Age in chimpanzees and gorillas are translated onto humans to control for cross-species variation in the pace of development and aging. Accordingly, survival rates are shifted in humans relative to great apes after cross-species age alignments. (C) Age alignments across humans, gorillas, orangutans, and chimpanzees.

### Individual Variation in Cross-Species Age Alignments

There is individual variation in the pace of development and aging. We collected time points across diverse human populations, sampled across Africa, Europe, Asia, and the Americas. We considered behavioral milestones, menarche, and age of peak births (populations: n=4, 47, 156 respectively; 29–31; Fig. 5). Variation in these biological and behavioral traits generally increase with age (Fig. 5C-F). The coefficient of variation (CV), which is calculated for time points from age of peak births, menarche, and behavioral milestones, extends up to 0.3 (5%-95%CI: 0.01-0.13). We consider this coefficient of variation captures plausible variation in extrapolated age alignments in humans and to be used as a guide in extrapolating ages across species.

We tested for sex differences in species for which sufficient time points were available for comparison. The pace of development and aging is similar across the sexes (Fig. 5A) in humans (n=134), chimpanzees (n=28), and gorillas (n=26). Many biological time points lie close to a y=x regression, which shows similarity in the pace of development across males and females. This is the case in humans (slope: 1.01 SE:0.003; y=1.01*log(x)-0.02; R2=0.99), chimpanzees (slope: 0.94; SE:0.033; y=0.94*log(x)+0.2, R2=0.97), and gorillas (slope: 0.99; SE: 0.1560; y=0.99*log(x)+0.072; R2=0.60. There is no obvious pattern where one sex takes longer to develop relative to the other across these three species. Notably, not all time points lie on a y=x regression (Fig. 5A). Some biological pathways occur for an unusually long time in one sex versus the other as is the case for body growth trajectories, which are extended in male gorillas (Fig. 5A; see sphere). Although some time points deviate from others across the sexes, males and females proceed through similar biological and behavioral trajectories.

### Testing Accuracy of Cross-Species Age Alignments

We tested the accuracy of our model in equating corresponding ages across species. We used maturational states of humans and great apes close to birth because of the availability of samples near birth. According to the model, a human in their first year after birth roughly equates to great apes in their first year of life (Fig. 5A). We considered cranial sutures (i.e., fontanelles) and cortical transcriptional profiles of humans and great apes close to birth (Fig. 6).

We aligned ages based on transcriptional profiles from the frontal cortex of 38 humans and 2 chimpanzees (Fig. 6B and C; 32). We correlated the log-transformed RPKM values of chimpanzees at days 0-1 post-birth with homologous genes in humans (n=12,557) ranging in age from close to birth to approximately 62 years of age. Only genes with a minimum expression were considered in these analyses (log10 (RPKM) >0.5). We found that the transcriptional profiles of chimpanzees around birth (Days 0-1 post-birth) most strongly correlate with humans at 1 year of age (Fig. 6C), highlighting the similarity in maturational state of humans and chimpanzees within the first year of life.

Next, we aligned ages based on the fontanelle maturity in humans and great apes. Human posterior and anterior fontanelles close at around 1 to 3 years (Fig. 6D). At birth, the anterior and posterior fontanelles extend across the medial to lateral axes of great apes and resemble that of human newborns (Fig. S14). The pattern of anterior and posterior cranial fontanelles of gorillas, orangutans, and chimpanzees near birth resembles humans that are also within their first year of life. Collectively, these observations suggest that great apes and humans are similar in degree of maturity in their first year of life, which agrees with the results from our model.

### Extended Duration of Biological Programs in Humans

Although the model accounts for a significant percentage of the variance in translated ages, some biological processes (e.g., carpal bone ossification, brain growth, locomotor behavior) deviate from others (Fig. 7; 33). Carpal ossification is one example where the timing of ossification is accelerated in great apes. We quantified ossified carpal numbers throughout development in humans, and some great apes. At birth, carpal numbers are generally higher in great apes than in humans, but these species differences are not significantly different (ANOVA: F=1.8; p=0.197; n=17; Fig. 7). Carpal bone ossification occurs at much faster pace in great apes (i.e., orangutans, and gorillas) than in humans. Ossified carpal bone numbers reach adult levels much earlier in orangutans (i.e., 1.5 years) and chimpanzees (i.e., 4.7 years) than they do in humans (11 years; Fig. 7). There are some sex differences in the rate of carpal ossification but both sexes reach adult numbers around 10 to 14 years of age (Fig. S16).

We classified each time point (i.e., brain growth, body growth, carpal ossification, organoid, cortical growth, locomotor development) and tested for deviations in the timing of biological pathways (Table S1, S9; Fig. 3, Fig. S1). The addition of body growth, brain growth, life history, structural aging, facial structure development, and locomotor forelimb development as factors in the model accounted for a significant percentage of the variance (Fig. 3). Therefore, there are significant heterochronies across species.

### Evolution of Lifespan in Human and Non-Human Primates

We evaluated whether the human lifespan is extended relative to great apes after cross-species age alignments (Fig. 8). Our translating time model captures time points up to 67 years of age in humans, which corresponds to 46 years of age in chimpanzees, and their equivalent in other primate species. We calculated the relative number of individuals (e.g., 90%, 80%, 70%) surviving up to a specific age (Fig. 8). Survival rates are extended in humans relative to great apes whether we align ages based on chronological ages (Fig. 8A) or map age in great apes onto human age (Fig. 8B). The extension in the human lifespan relative to great apes does overlap with some human populations but is clearly shifted relative to most human populations. These plots demonstrate that the human lifespan is extended relative to studied great apes.

## Discussion

We generated cross-species age alignments across humans and other primates. We identified which biological programs are conserved and which have been modified in humans. The inclusion of time points from diverse populations captures individual variation. One important finding from the present study is that the human lifespan is extended relative to great apes after cross-species age alignments.

### Age Alignment Across Humans and Non-Human Primates

This work expands on a long-term project called Translating Time (www.translatingtime.org), which relied on abrupt transformations to align ages during prenatal and early postnatal development in humans and model systems (18, 34). Here, we aligned biological pathways from abrupt and gradual changes in transcriptional, structural, and behavioral variation in order to find comparable ages across the lifespan of human and non-human primates. We tested several machine learning techniques to find models best suited to generate cross-species age alignments. This integrative approach expanded our dataset by an order of magnitude relative to past studies (34, 35).

Our model aligns ages across species. Early in development, corresponding ages are similar across humans and great apes but they gradually diverge with age (Fig. 8). For instance, a human in their first year of age equates to a gorilla, orangutan, bonobo, and chimpanzee at roughly similar ages but cross-species differences in the pace of development and aging become salient with age (Fig. 8).

Capturing information across diverse populations can be used to quantify variation in translated ages. Environmental factors also impact the pace of development and aging with noticeable differences between captive and wild populations. We captured time points from diverse human populations, and from great apes that were mostly held in captivity rather than from the wild, and the results from our study mostly apply to captive primates. One notable observation is that the pace of development and aging are similar, but a subset of time points deviate from most others within each species (Fig. 5). Our dataset only captures age ranges up to mid-teens in great apes. We have yet to evaluate potential variation in the pace of aging in these species and investigate these sex differences. We plan to do so in future studies.

### Modifications in Carpal Ossification

We found that some time points were protracted relative to other time points in some species. In particular, the rate of carpal bone ossification is accelerated in great apes relative to humans and is likely linked to species differences in locomotion. Knuckle walking, which is specific to great apes, and our study shows that species differences in these forelimb locomotor adaptations are linked to carpal bone ossification acceleration.

### Old Age as a Distinctively Human Feature

We found that the human lifespan is unusually extended compared to great apes, and which is a pattern that holds across populations. It is rare that great apes live beyond 50 years of age, though one chimpanzee was recently estimated to live up to 68 years of age (36–39). Lifespans are malleable and enhanced care of great apes could lengthen lifespans. Nevertheless, we suggest that the extension in human lifespan explains species differences in biological pathways in old age. The shorter lifespan of great apes relative to humans may explain the difficulties in observing plaques, tangles, and brain atrophy in great apes (39–41). Extending great ape lifespan may reveal biological processes, including pathologies, that are currently thought to be unique to humans.

In old age, humans and great apes suffer from similar but also non-overlapping diseases, which may contribute to species differences in lifespan (41–44). Chimpanzees and humans suffer from heart diseases, but chimpanzees are more likely to suffer from interstitial myocardial fibrosis whereas humans are more likely to suffer from coronary-artery atherosclerosis (43). Moreover, humans are more likely to suffer from cancer relative to chimpanzees (41, 45), but chimpanzees are more likely to die of viruses and bacterial infections than humans. Information on disease incidence in great apes are from captive records. Disease incidence is likely heavily impacted by environmental factors, including stress and diet, and the relative disease incidence likely varies between wild and captive populations of great apes.

### Translating Time in Great Apes Enhances Conservation Efforts

The present study provides translational tools to find equivalent ages across the lifespan of humans and great apes (e.g., orangutans, gorillas). It is expected this work will enhance our ability to detect abnormalities at early stages, and improve timeliness of interventions in both humans and in non-human primates. The ability to track development in critically endangered species such as gorillas and orangutans, for which data on biological timelines are sparse, can improve treatment and increase population size of these threatened species. The results from the present study can now be used as a baseline with which to detect normal developmental timelines in studied great apes.

## Conclusions

Our work builds a resource to align ages across humans, great apes, and monkeys. We identified which biological programs are conserved and which have become modified over the course of development and aging. Capturing these basic parameters across human and nonhuman primates can be used to enhance tracking capabilities in humans as well as in great apes.

## Materials and Methods

We gathered time points from abrupt and gradual changes in transcriptional, anatomy, and behavior across 9 species (Figs. 1 and S1) and from diverse human populations (see SI Appendix; Figs. 1 and 5; Table S1). Data was imputed to generate an event scale (see SI Appendix). Statistics were performed with the programming language R.

### Behavioral and Structural Variation for Age Alignments

We used bone radiographs, some of which were provided by the North Carolina Zoo. These data were used retrospectively, collected for purposes other than this study, and were approved for use by the North Carolina Zoo IACUC committee. We used these and other images to track carpal ossification maturation (Fig. 7E-I). We quantified the number of discernable ossified structures in the wrist (Figs. 7B and S16).

We extracted time points from peaks and plateaus in growth trajectories (Figs. 1–3). We fit non-linear regressions through the data to extract the age at which individuals reach specific percentages of adult volumes (Figs. S8–S10). We also fit non-linear regressions to capture the age at which peaks in specific biological processes occur across different species (e.g., age of peak births; Fig. S13). Some data were extrapolated from the Web Plot digitizer. In some cases, data points may have been obscured by others on plots or data were collected from regressions. Therefore, time points collected may vary slightly from that reported in the original study.

### Transcriptional Variation for Age Alignments

We trained 6 machine learning models to predict corresponding ages from normalized gene expression from individuals at different ages (20, 32, 46–48). We also used human and gorilla organoids to generate cross-species age alignments (Fig. 5; see SI Appendix). These data are from publicly available datasets (32, 47, 48).

## Acknowledgments

We thank Dr. Melissa Harrington for her support and Emily Lynch for access to radiographs. Some data are from the Allen Institute and the Brainspan Atlas of the developing human brain (http://www.brainspan.org; http://developingmouse.brain-map.org). The acquisition of some data were supported by the National Institutes of Health (NIH) Contract HHSN-271-2008-00 047-C. Some scans are from the Digital Morphology Museum, Kyoto University Primate Research Institute (KUPRI). This research made use of the Primate Aging Database (https://primatedatabase.org/), which is an initiative of the National Institute on Aging. We thank Christine Vincent for quality control and Javier Lazzaro for drawings, some of which were created with BioRender.com.

## Supplementary material

We discuss how we filtered the genes for the application of machine learning models, how we imputed the data to generate the event scale, and the different sources that make up the dataset.

### Variation across human populations

We collected time points from individuals in the wild and in captivity. We used menarche, age of peak births, and survival rates across diverse populations. We included survival rates from multiple sources from captive and wild populations. Survival rates were computed by quantifying the number of individuals at a particular age relative to individuals present at or close to birth. We only included survival rates from populations with a relative clear decline in the percentage of individuals (1, 2).

### RNA metrics for machine learning analyses

We filtered genes with low average expression. Poorly expressed genes are typically noisier in their expression than highly expressed genes. We aimed to use a roughly similar number of genes sampled across datasets (~10,000 to ~13,000 genes). We performed analyses across humans and chimpanzees versus other nonhuman primates. We performed additional analysis where we extrapolated ages across humans and chimpanzees from normalized gene expression from the prefrontal cortex and included them in the dataset (Table S1). For some tested models, there were missing values in resampled performance measures. We kept outputs of all six models across tested datasets to readily compare their performance across datasets.

### Imputation for the model generation

We fit a linear model to translate ages. We imputed time points because data were not systematically available across all 8 studied species. Orangutans and gibbons were amongst the species for which data are sparse whereas data for humans and chimpanzees’ data are relatively complete (Fig. S1). We imputed missing data with imputation by linear regression through a prediction method implemented with a Monte Carlo simulation. We used the midas touch method, which imputes univariate missing data through predictive mean matching. We selected the imputed dataset with the highest correlation coefficients across individuals. Specifically, we evaluated the lowest correlation coefficient from each dataset, and we selected the dataset with the highest minimum correlation across individuals. We ordered these time points and subtracted each time point (averaged across species) by the minimum age and divided these values by the difference between the maximum and minimum average age. The ordering of time points varied from 0 to 1 with early time points assigned a score close to 0 and later time points assigned a score close to 1. We collected the average time points across species with the weighted average proportional to the amount of missing data so that species with relatively fewer time points contributed less to the events scale than species with more complete data.

### Data source

Time points are collected from multiple sources. Some time points were collected from multiple individuals, but other age alignments were collected from a single individual. Data on great apes are sparse, especially for highly endangered gibbons and orangutans (Fig. S1). We chose to maximize sample size even though sex and environmental conditions (i.e., captive versus wild) were not systematically known for every time point. We tagged time points from males versus females when this information was known (Fig. 5A and B). We did not classify individuals by sex for prenatal and early postnatal time points because information about sex is frequently lacking from these developmental studies. We used multiple metrics where possible as well as large samples to overcome potential inconsistencies (Table S1). Despite these caveats, our approach resulted in an unprecedentedly large dataset and enables the translation of ages for highly endangered great ape species.

**Fig. S1.**
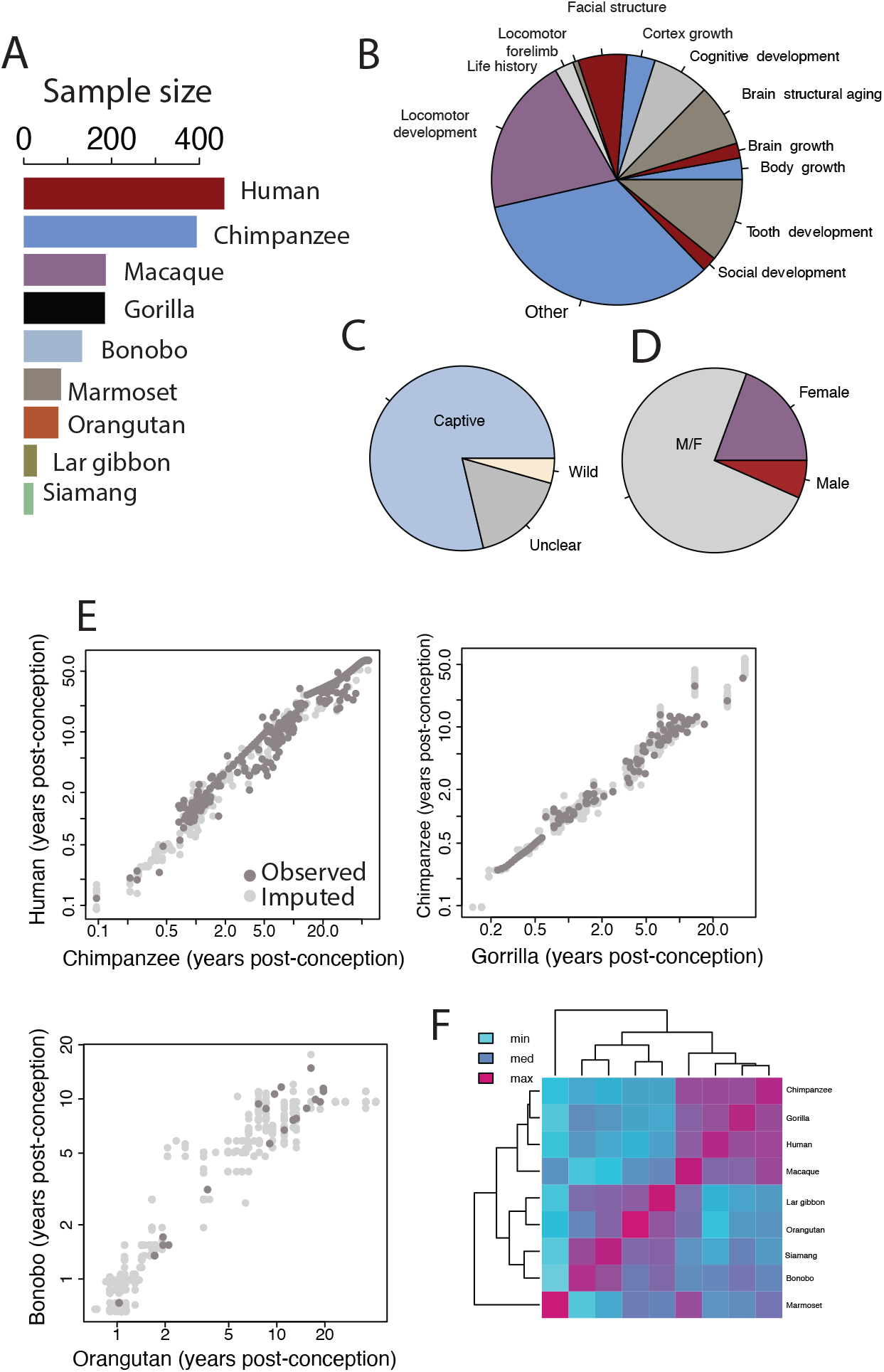
(A) A barplot shows the number of time points collected for each species. More data were collected for humans and chimpanzees than for other species (e.g., gibbons, orangutans). We combined information for different subspecies of orangutans and gorillas due to the paucity of data available for orangutans and gorillas. (B) The data is largely composed of structural and behavioral time variation. (C) Most of the data are from captive rather than wild primate populations, and (D) most of the data is either averaged by sex or of unknown sex. (E) We imputed time points to generate an event scale. Examples are shown for humans, chimpanzees, and gorillas. Correlation coefficients from log-transformed time points for different species are higher for gorillas, chimpanzees, macaques, and humans (pink) than they are for gibbons and orangutans. That is, gibbons and orangutans correlate least with other species, and are those with the least amount of time points.

**Fig. S2.**
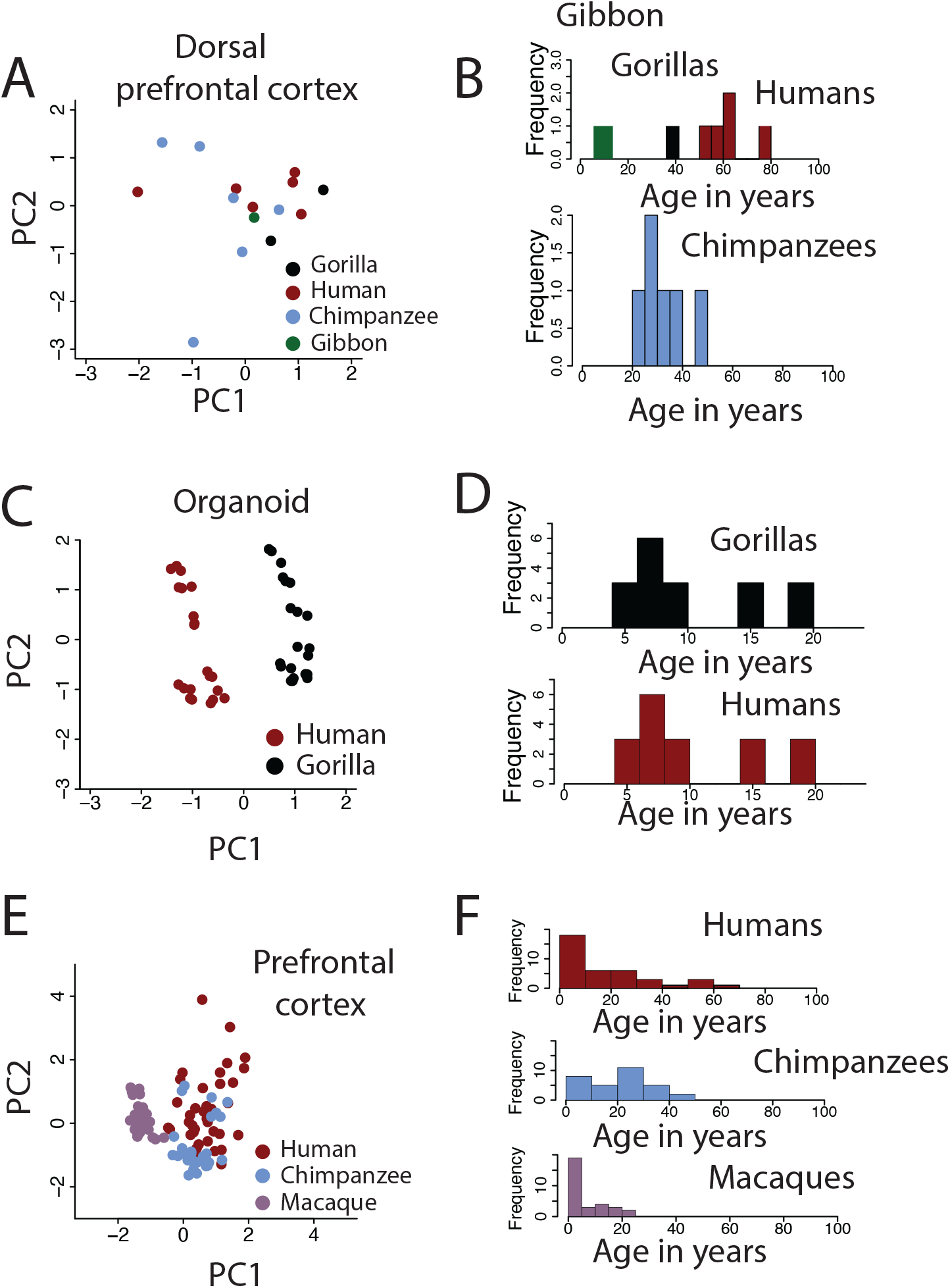
We used RNA sequencing datasets from multiple sources and from multiple species to generate cross-species age alignments ((3); A-B), ((4); C-D), and ((5); E-F). (A, C, E) We performed a principal component analysis of log10-transformed normalized gene expression. (B, D, F) We also include histograms to include the age ranges for each dataset. Age here is expressed in years after birth of individuals. We used these data to test which machine learning models are best suited to align ages across human and non-human primate species.

**Fig. S3.**
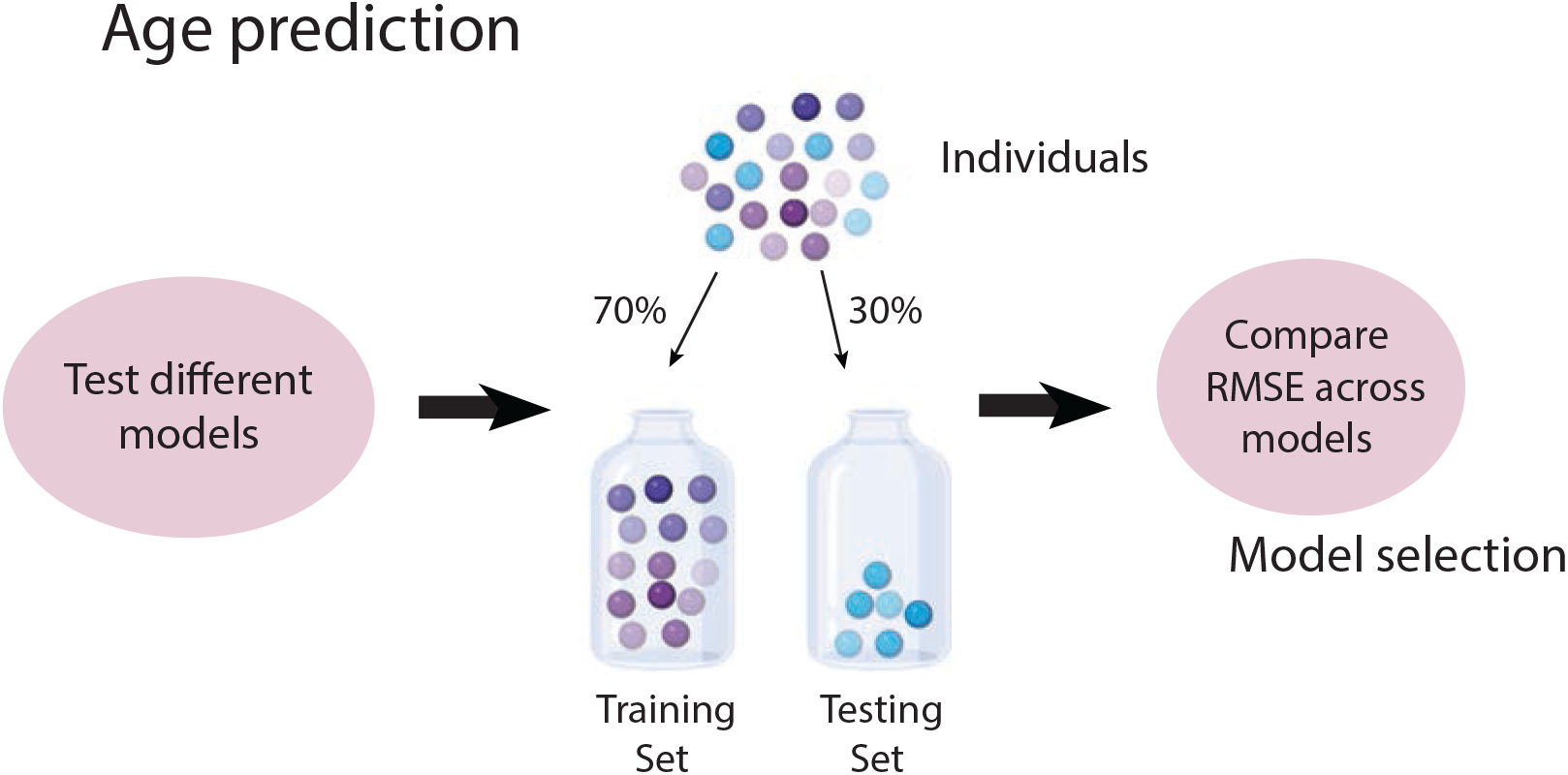
Overview of the approach used to generate cross-species age alignments from machine learning models. We first partitioned the data into a training and a testing set. The training set consisted of roughly 70% of the data, and the testing set consisted of the remaining 30% of the data. We trained 6 different models (e.g., random forest) to predict ages, and we checked the accuracy of these predictions. We use the root mean square error (RMSE) to select the best suited model. RMSE values measure the difference between actual and predicted values. We then generated cross-species age alignments by inputting data from one species model into these trained models.

**Fig. S4.**
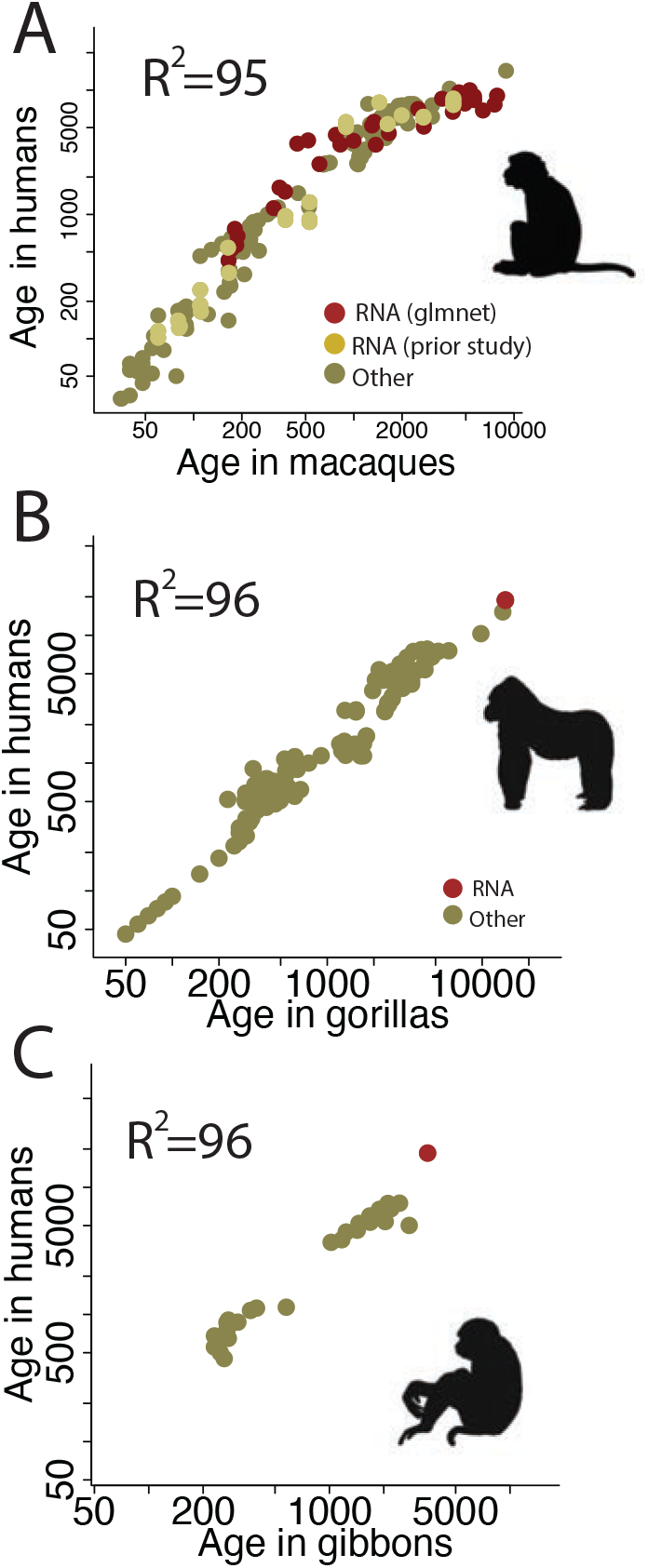
We compared corresponding ages extracted from machine learning models with other metrics to assess the accuracy of translated ages. (A) Corresponding time points extracted from machine learning models, other statistical procedures from a past study (6), as well as other metrics (mostly structural and behavioral metrics) yield similar results. We fit a linear regression through the log-transformed values expressed in days after conception, which accounts for a significant and high percentage of the variance (95.6%), and this analysis is modified from a recent study (7). (B) We also extracted time points from machine learning models across humans and gorillas. We found that time points from machine learning models align with other metrics. The linear regression across these data accounts for a significant and high percentage of the variance (95.7%). (C) We also used machine learning models to translate ages in gibbons. The time point extracted from machine learning models in gibbons appears to deviate from other time points.

**Fig. S5.**
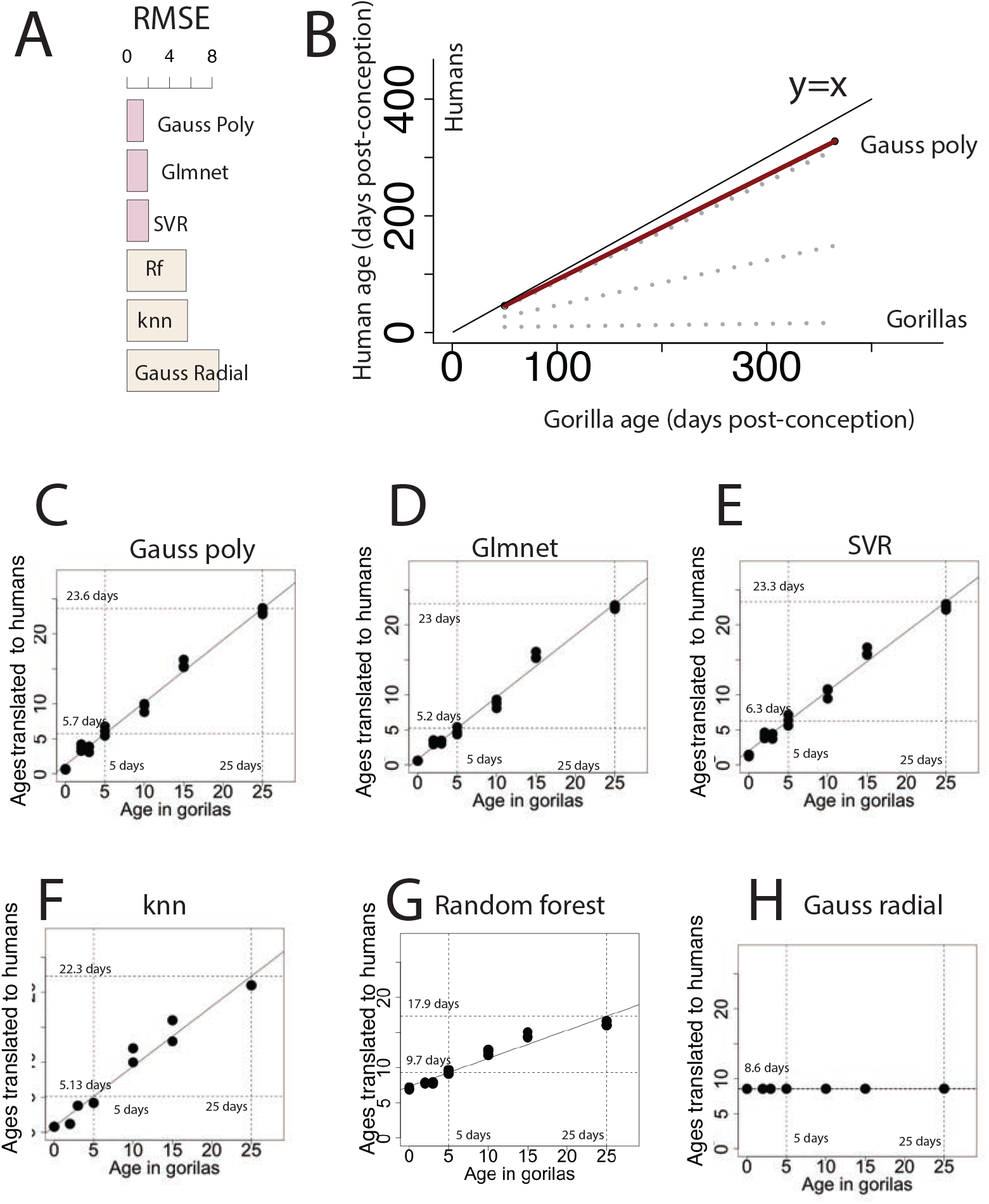
We tested 6 machine learning models to generate cross-species age alignments from human and gorilla organoids. (A) The glmnet R package, gauss poly, and SVR were considered the best models to translate ages because they produced the lowest RMSE values. Different models produced a range of corresponding ages across the two species. (B) We compared how different models yielded different translated ages in humans and gorillas. We generated a regression through human and gorilla time points from organoids and used this regression to extend ages across prenatal stages in humans and gorillas. (B) We included a y=x regression as a baseline to detect whether translated ages are similar in absolute days across these two species. That is, regressions that lie close to y=x mean that corresponding ages are similar across the two species. We selected translated ages produced by the gauss poly model. (CH) Translated ages according to different models. The gauss poly (C), glmnet (D), and SVR model (E) were considered the best models based on RMSE values.

**Fig. S6.**
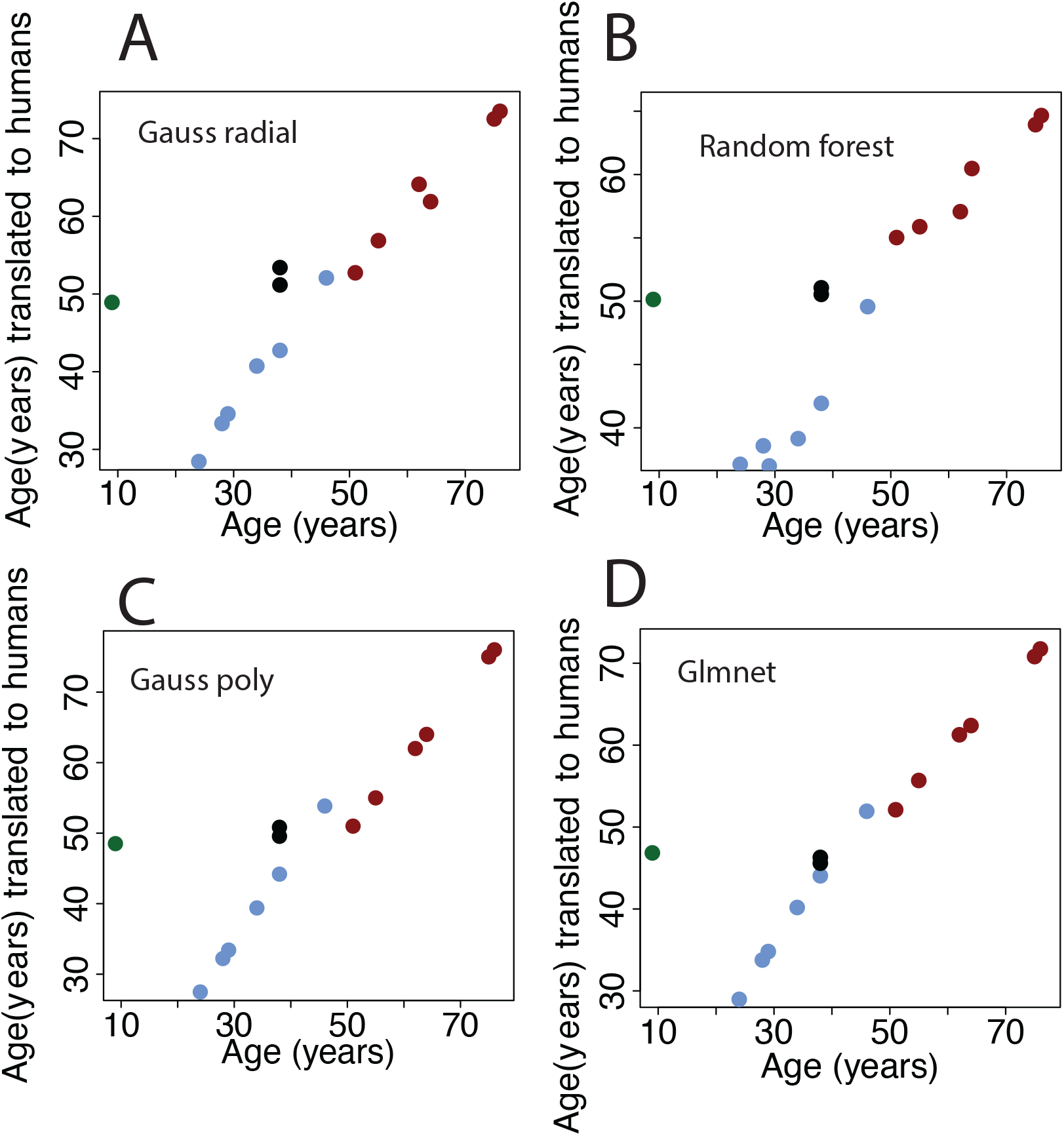
We used different models to compare age alignments from normalized gene expression extracted from the frontal cortex of humans, chimpanzees, gibbons, and gorillas. (A-C) We considered how these models predict age in humans for which age is already known.

**Fig. S7.**
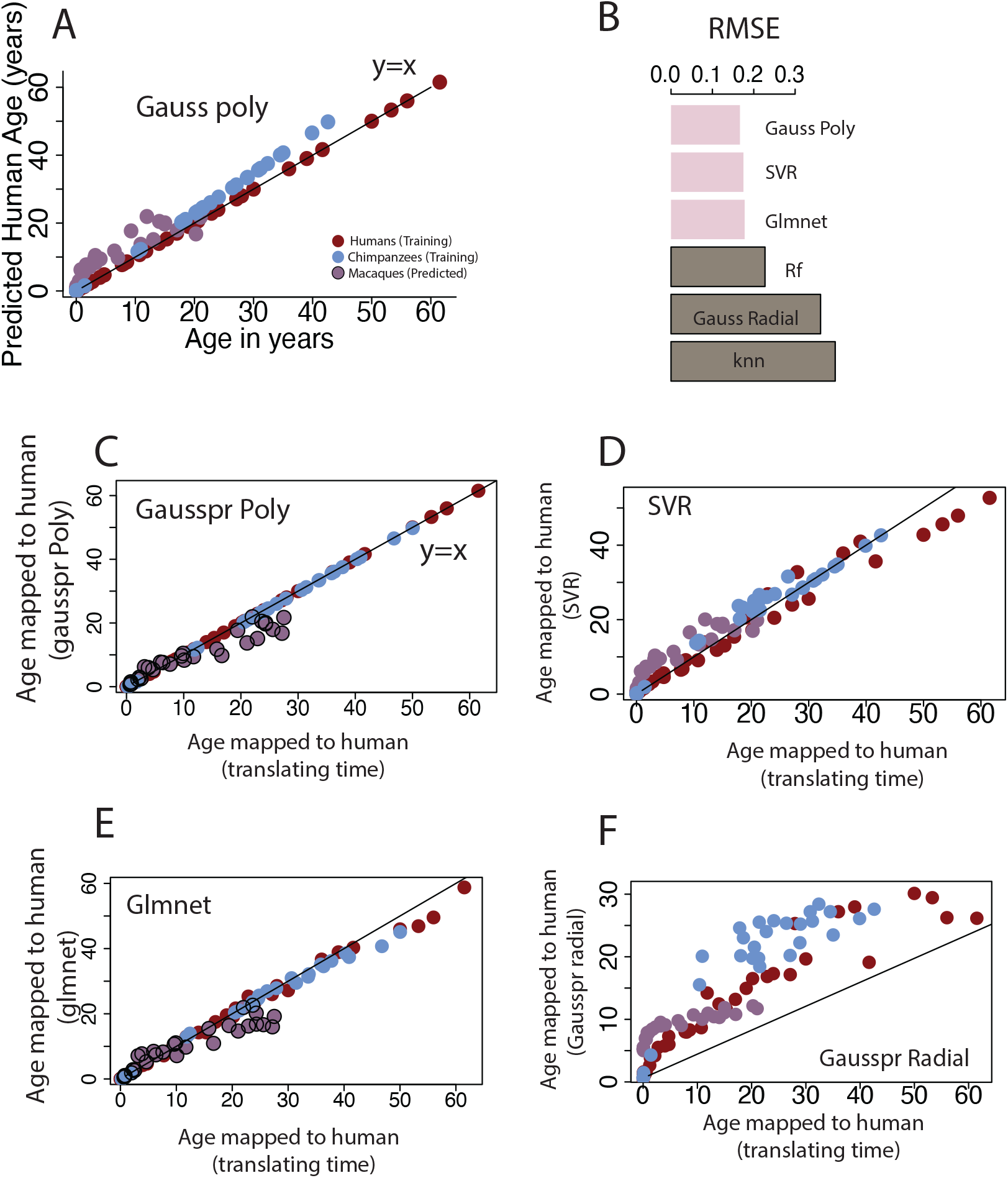
(A) We used machine learning models to find corresponding ages between species. Here, we used one of these models to predict age from normalized gene expression from human, macaque, and chimpanzee frontal cortex. (B) Model selection was based on RMSE values (B), which compares the difference between actual and predicted ages. A low RMSE indicates good model performance. (C-E) We compared translated ages based on our machine learning models to our previous work collected (7, 8) to ensure the concordance of age alignments based on different methods. We mapped the age of chimpanzee and macaques onto humans (on the y axis) and we mapped ages of chimpanzees and macaques onto humans based on past work (x.axis; (7, 8)). Time points that lie in close proximity to the y=x regression demonstrate strong concordance across methods. Translated ages extrapolated from the gauss poly generated best prediction accuracies.

**Fig. S8.**
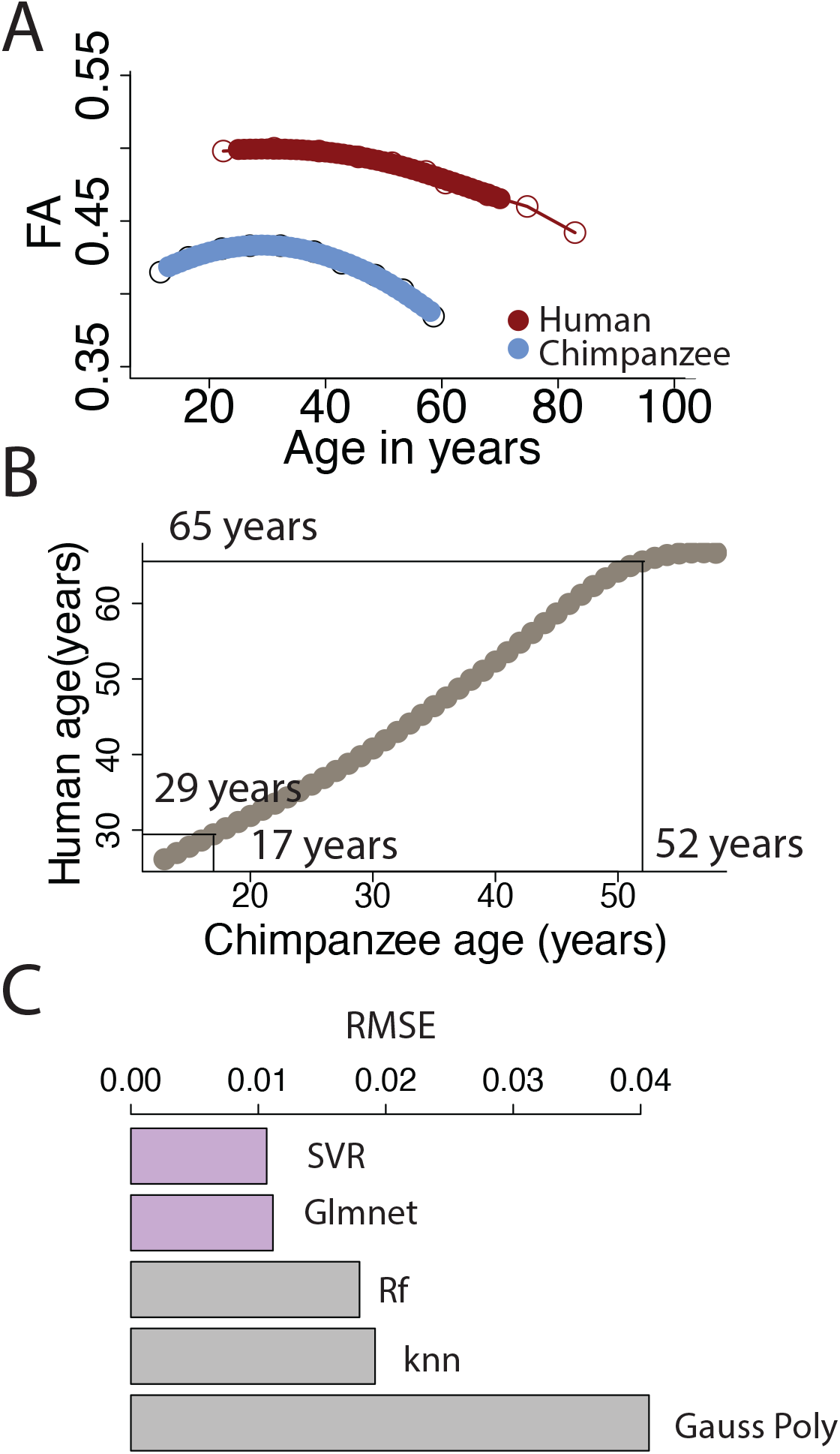
We used fractional anisotropy, radial and mean diffusivity to align ages from machine learning models. (A) Fractional anisotropy varies similarly with age in humans and in chimpanzees. (B) We applied different machine learning models to extract corresponding ages from these metrics. (C) The glmnet model produced the lowest RMSE scores. We therefore used the SVR model to align ages across the two species. Corresponding time points occur later in humans than in chimpanzees.

**Fig. S9.**
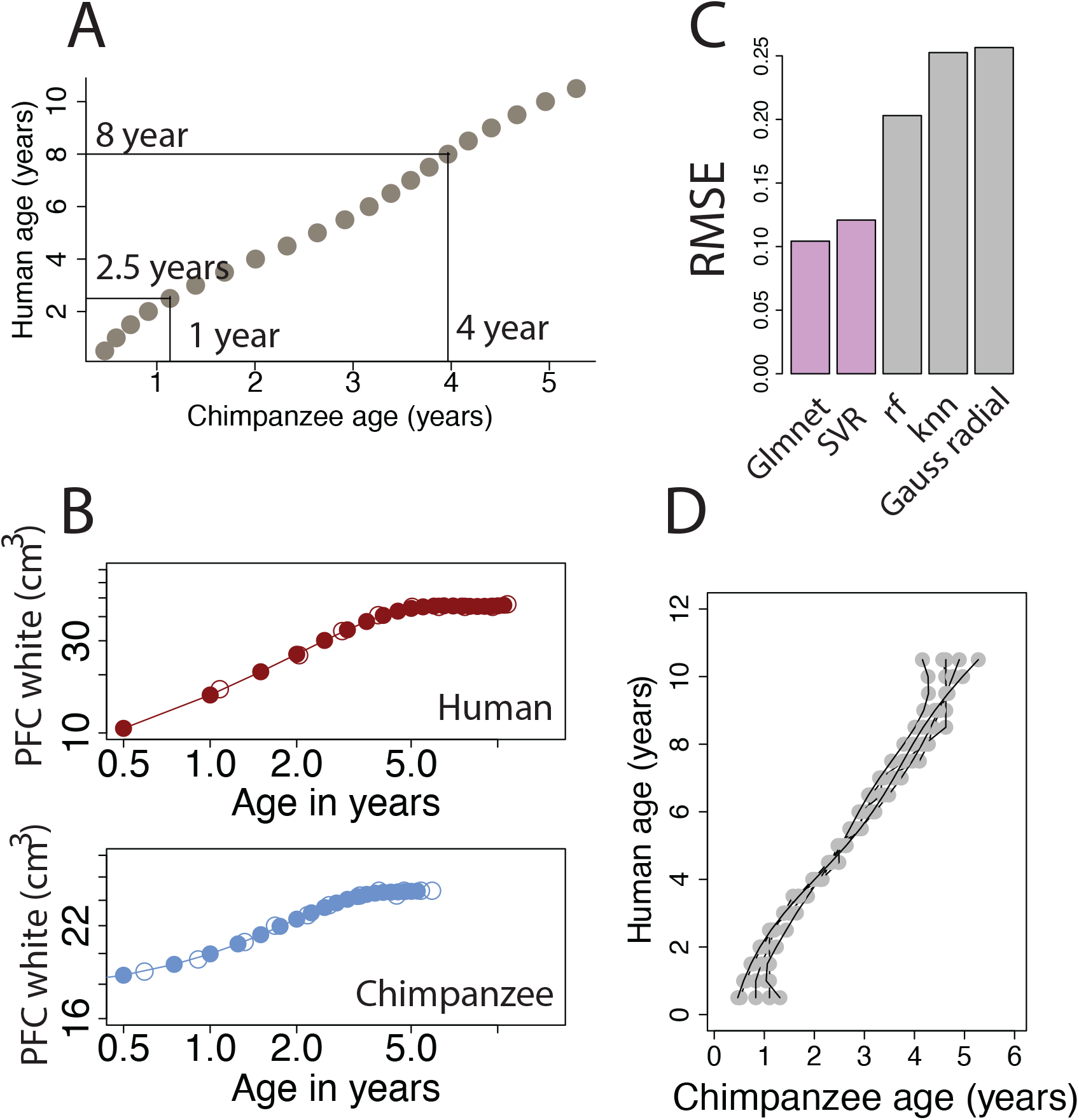
(A) Age alignments generated from the growth of frontal cortical areas and the corpus callosum in humans and in chimpanzees. (B) We used the prefrontal cortex white matter volumes from humans and chimpanzees of different ages as one of multiple metrics to align ages across these two species. We first trained the model to predict age within a species, and relied on RMSE values as a basis for model selection. (C) RMSE values showed that the glmnet model package produced the lowest RMSE values. We therefore selected the glmnet model to generate cross-species alignments. (D) We also include cross-species age alignments produced from different machine learning models for comparison. Corresponding time points occur later in humans than they do in chimpanzees.

**Fig. S10.**
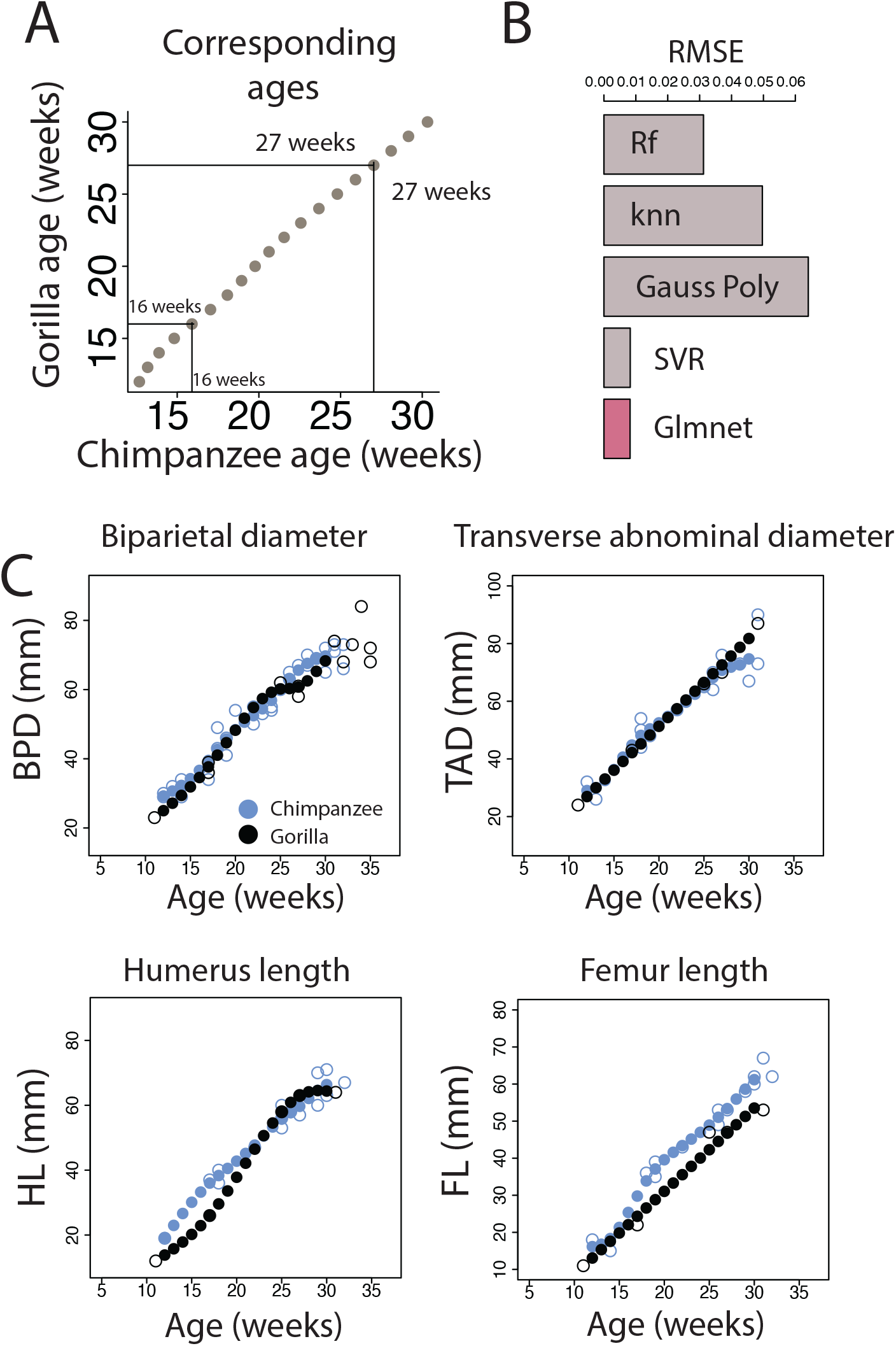
We aligned ages (A) from temporal growth metrics extracted from ultrasounds of fetal gorillas and chimpanzees. (B) We tested different machine learning models, and we selected the glmnet model based on RMSE values. Metrics used in the model include (C) include biparietal diameter (BPD), transverse abnormal diameter (TAD), humerus (HL), and femur length (FL). We fit smooth splines through the data in order to capture the same sample size across gorillas and chimpanzees. These analyses enable finding corresponding ages during fetal stages of development across great apes.

**Fig. S11.**
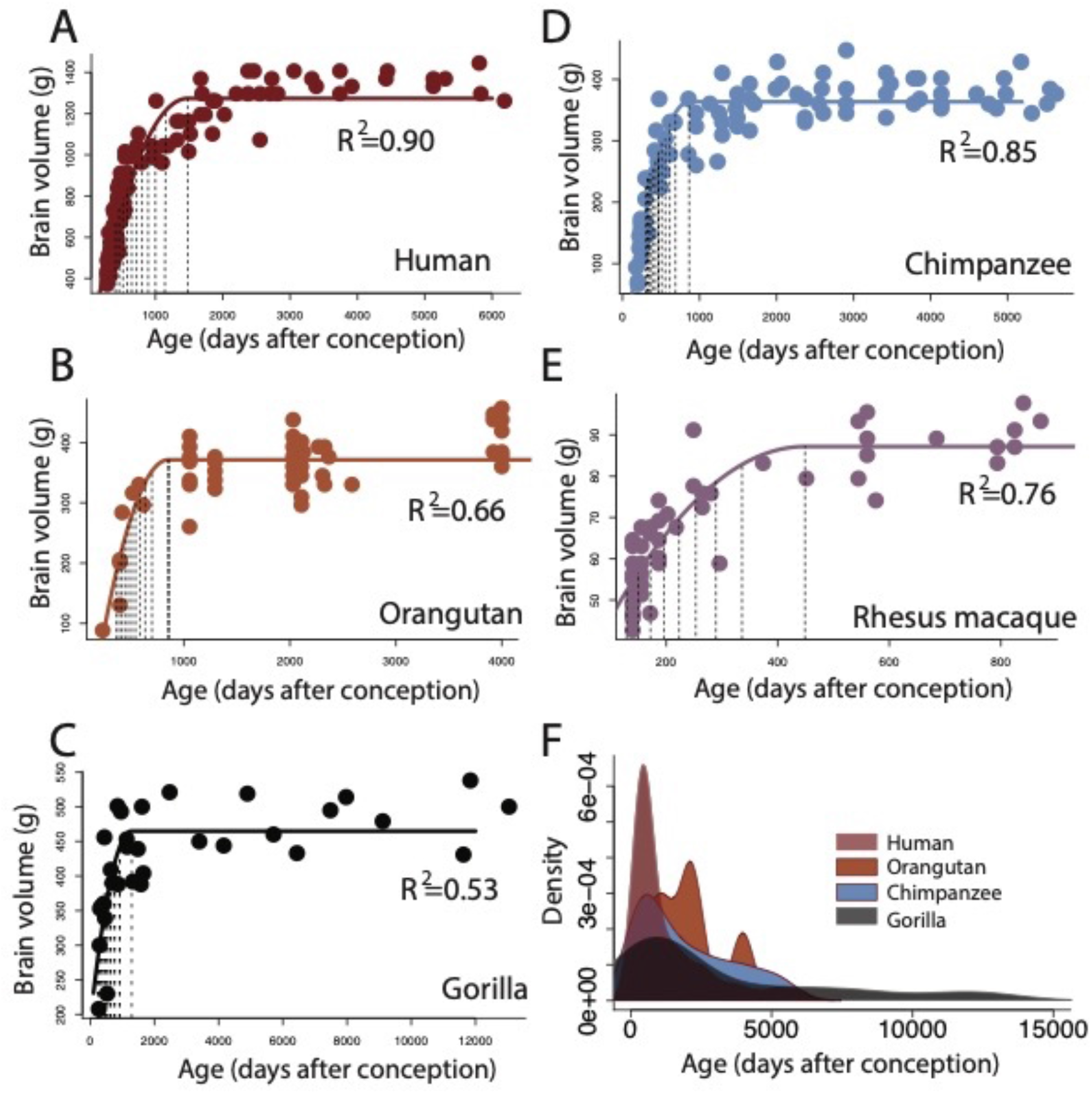
We extracted time points from brain growth trajectories in humans (A), orangutans (B), gorillas (C), chimpanzees (D), and rhesus macaques (E; (9)). We fit non-linear regressions (easynls, model=4) with age as the independent variable and brain volume (grams) as the dependent variable. We extrapolated the age at which the brain reaches adult volumes and percentages of adult volumes (vertical bars) from these non-linear regressions. These regressions capture when most of the brain ceases to grow but some, albeit small, growth may extend beyond these identified ages. Species differences in sample size may introduce variation in identified age of growth cessation across species. We, therefore, excluded time points that occur before 270 days after conception in humans and their corresponding ages in macaques (127 days after conception; (10)) to capture similar age ranges across species. This is because brain growth appears accelerated at prenatal stages relative to postnatal ages in humans and macaques. Fetal data was not available for all studied primate species but their inclusion in a subset of studied species could impact the output of the regression. (F) Kernel density plots show the age ranges for available data.

**Fig. S12.**
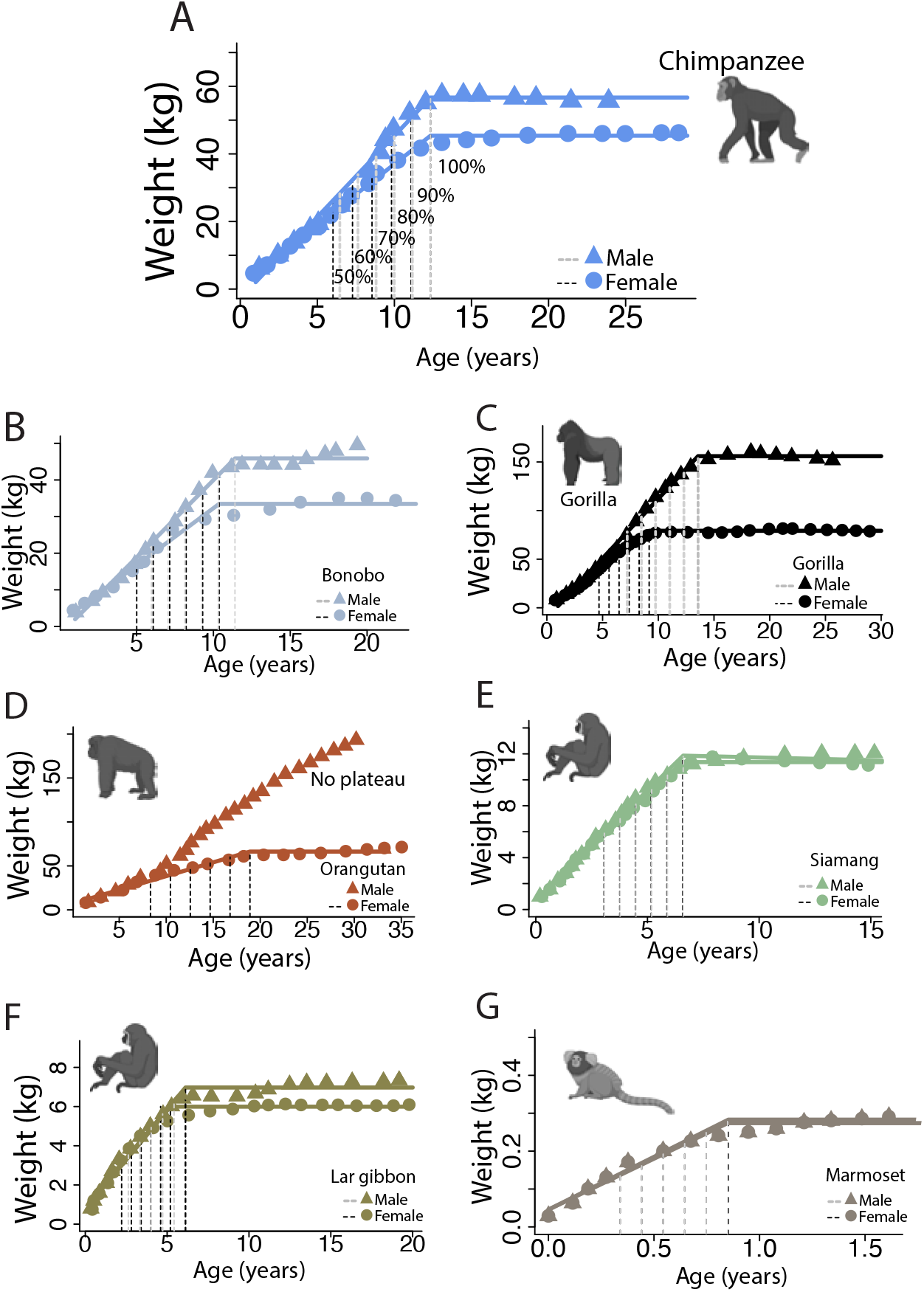
We include corresponding time points from body growth trajectories from great apes (A-D), lesser apes (E-F), and from monkeys (marmosets; G). Humans are also in the model. We fit a nonlinear regression (easynls, model=3) with weight versus age expressed in years. We fit regressions separately for males (triangles) and for females (circles). Sex differences are noticeable in some of the studied primates (e.g., gorillas, C; orangutans, D). We extracted ages at which the percentage of adult volume reaches a particular value (e.g., 100%, 90%, 80%) as a basis with which to equate corresponding ages across species. The dashed vertical lines represent time points extracted from these growth trajectories collected separately for males and females and are exemplified in chimpanzees (A). Not all groups reach adult weights. This is notably the case for male orangutans that grow throughout their life. We did not extrapolate time points for male orangutans. A greater age range is available for humans than for other studied non-human primates, which might skew the extrapolated time points. We therefore fit nonlinear regressions through weights for individuals from 2 to 34.5 years of age so that non-linear regressions are collected through similar age ranges across non-human primates. These data are from captive individuals.

**Fig. S13.**
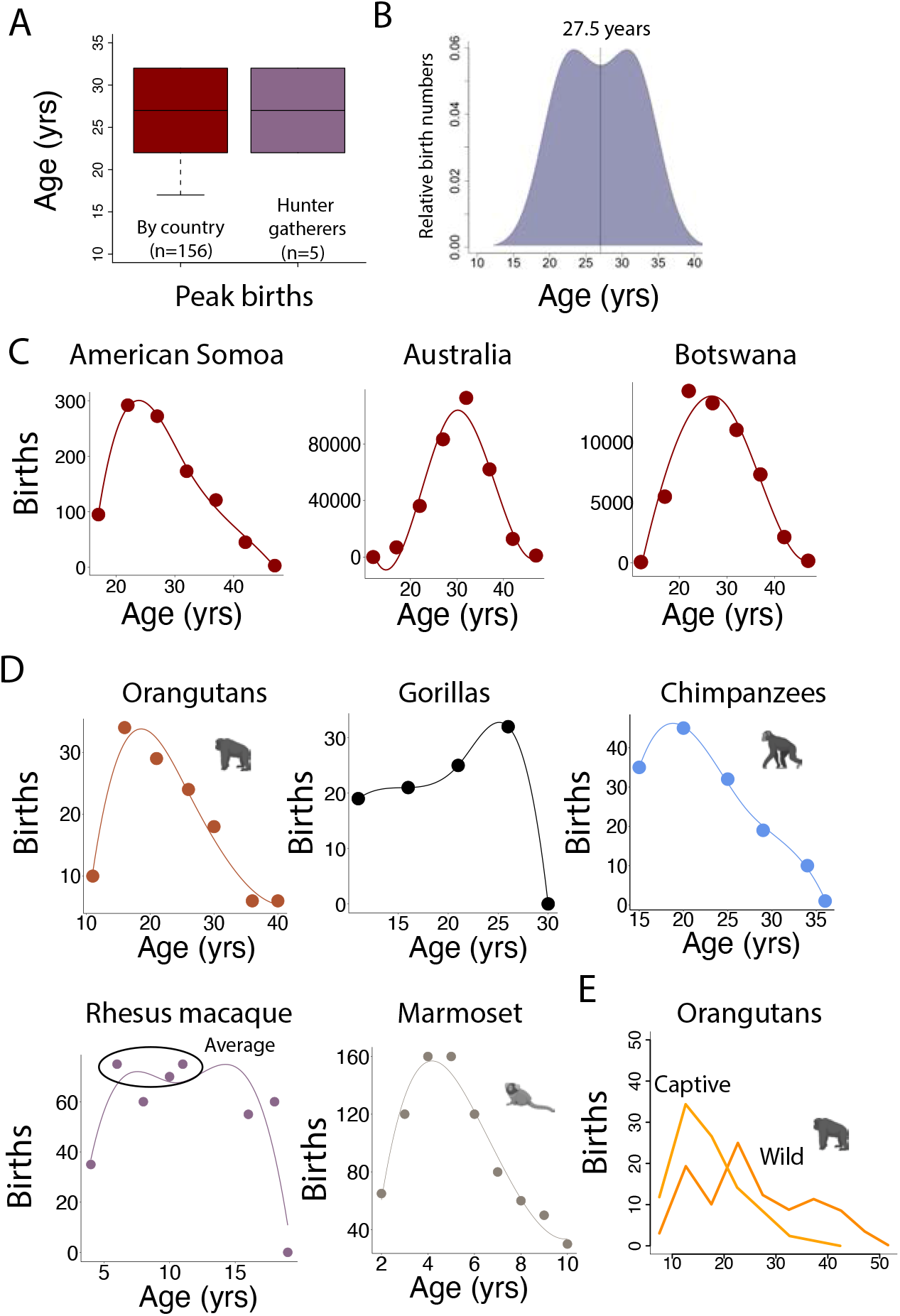
We considered individual variation in the age of peak births across human populations across 156 countries as well as from hunter-gatherers. We also include the age of peak births for some non-human primates (i.e., chimpanzees, gorillas, orangutans, macaques, marmosets). (A) We considered the age of peak births from hunter-gatherers (B; n=5), and from industrialized societies. (C) We include a few examples, which include birth rates from American Samoa, Australia, and Botswana, to exemplify variation across diverse human populations. The 95% confidence intervals in the age of peak births range between 22 to 32 years of age in humans with a similar median average age between hunter-gatherers and industrialized societies. (D) We obtained the age of peak births from great apes (e.g., orangutans, gorillas, chimpanzees) and monkeys (e.g., rhesus macaques, marmosets; (11)). The 95% confidence intervals in the age of peak births in non-human primates range between 4 (marmosets) to 25 years of age (gorillas). The y axis shows the mean number of births per female in a given interval of time and are shown as a percentage. (E) Age of peak births can vary dramatically based on environment as evident between captive and wild orangutans.

**Fig. S14.**
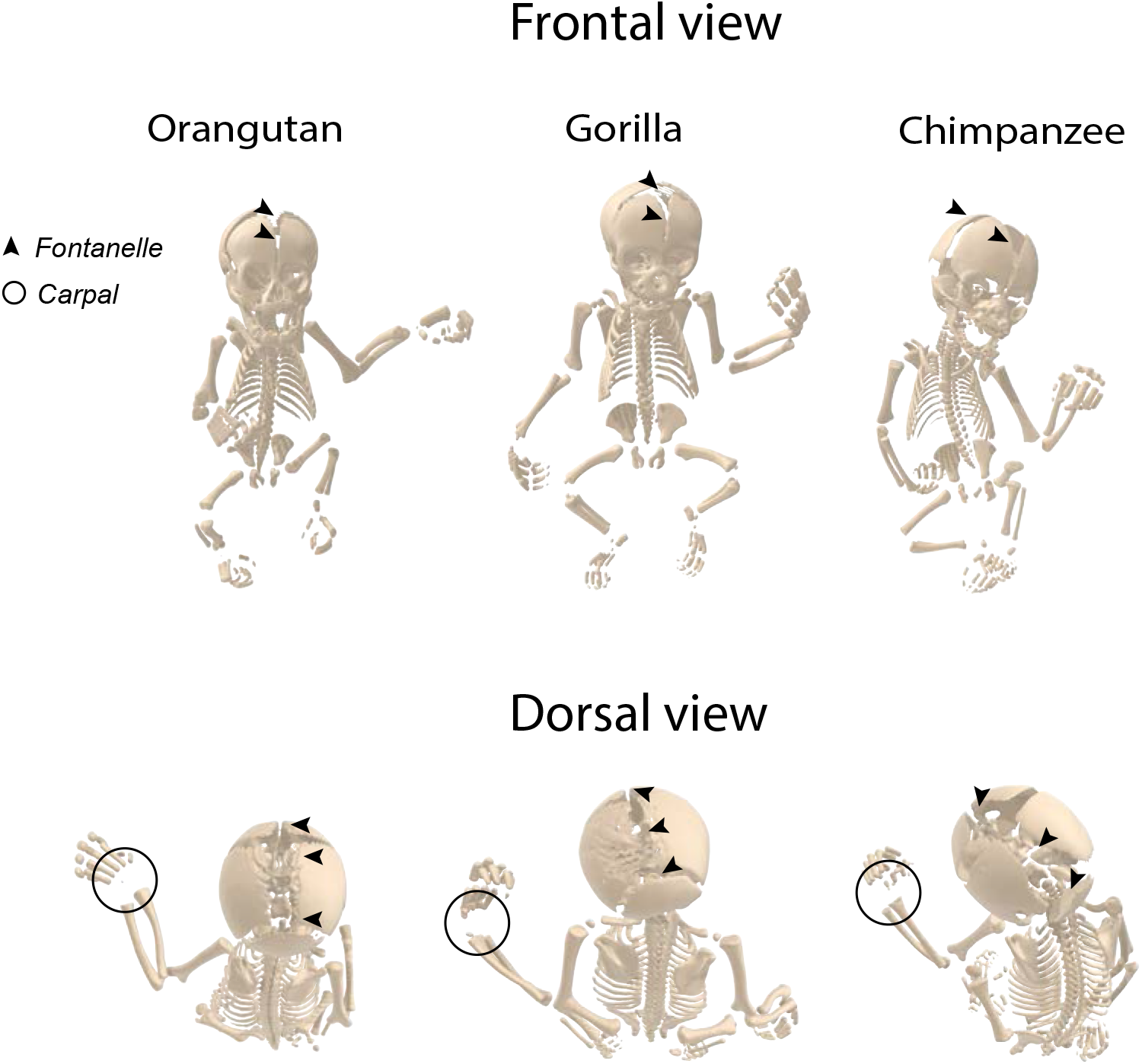
We used CT scans of a chimpanzee, gorilla, and an orangutan estimated to have died close to birth. We use the maturity of key phenotypic traits (e.g., fontanelles, carpal bone ossification) as a complimentary approach to align ages across humans and apes to align ages across birth. We considered the presence of fontanelles and ossified carpal bones in great apes and in humans to align ages and detect possible deviations in the timing of biological pathways. According to the scans of these individuals, fontanelles have yet to close in these specimens, and this is true of the gorilla, orangutan, and the chimpanzee. In addition, carpals have yet to be ossified. These scans are from the visible ape project (12).

**Fig. S15.**
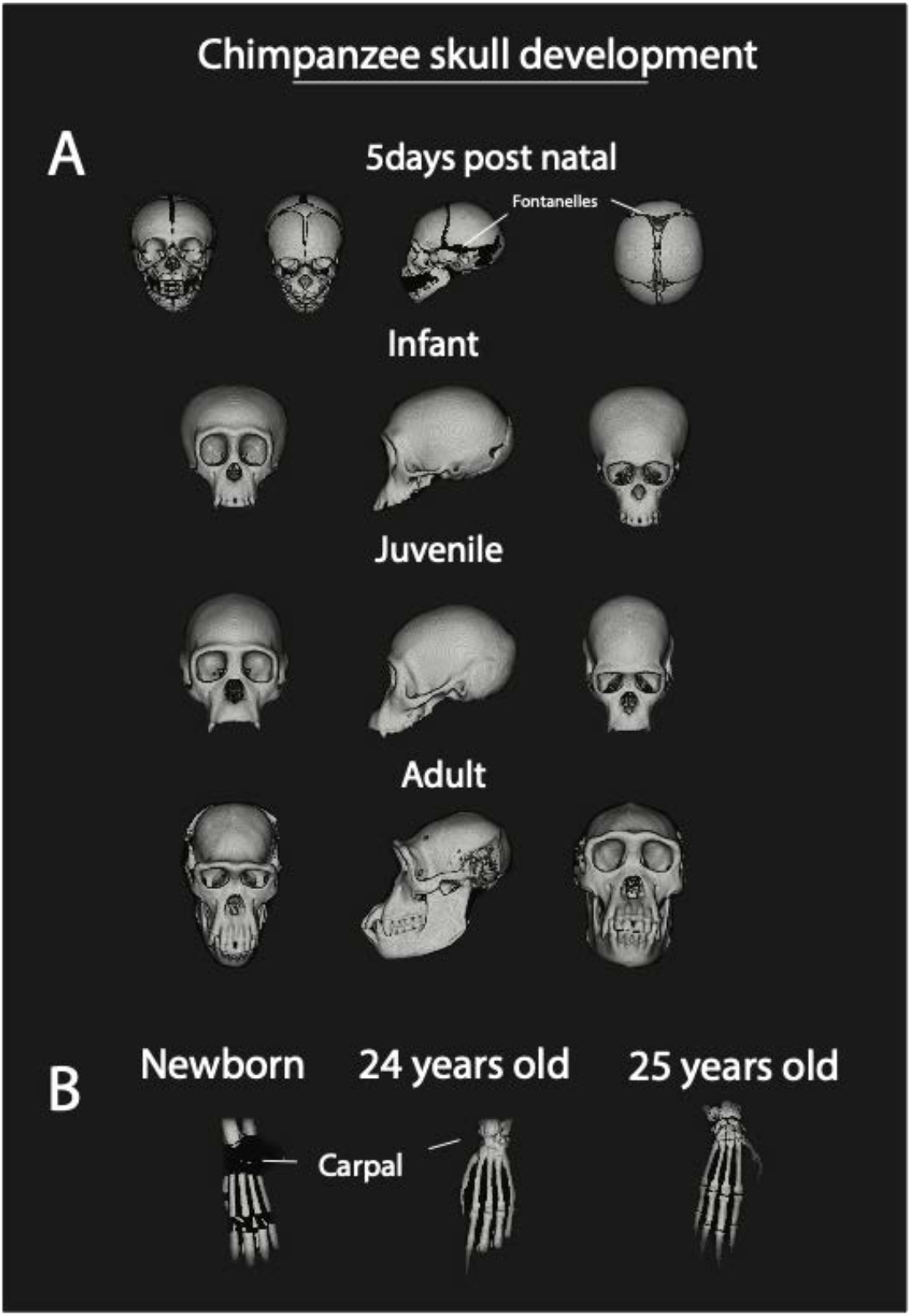
We considered skull (A) and wrist maturation (B) from CT scans of chimpanzees from the Digital Morphology Museum, Kyoto University Primate Research Institute (KUPRI). Fontanelles are evident in the 5-day old newborn chimpanzee but not in the infant. Infancy is typically defined as ranging between birth to about 3 years of age (8). According to these observations, fontanelle closure should occur somewhere between 0 to 3 years of age. Similarly, carpal bone ossification (B) extends postnatally. This is evident given that the 5-day old chimpanzee has many carpals that have yet to ossify.

**Fig. S16.**
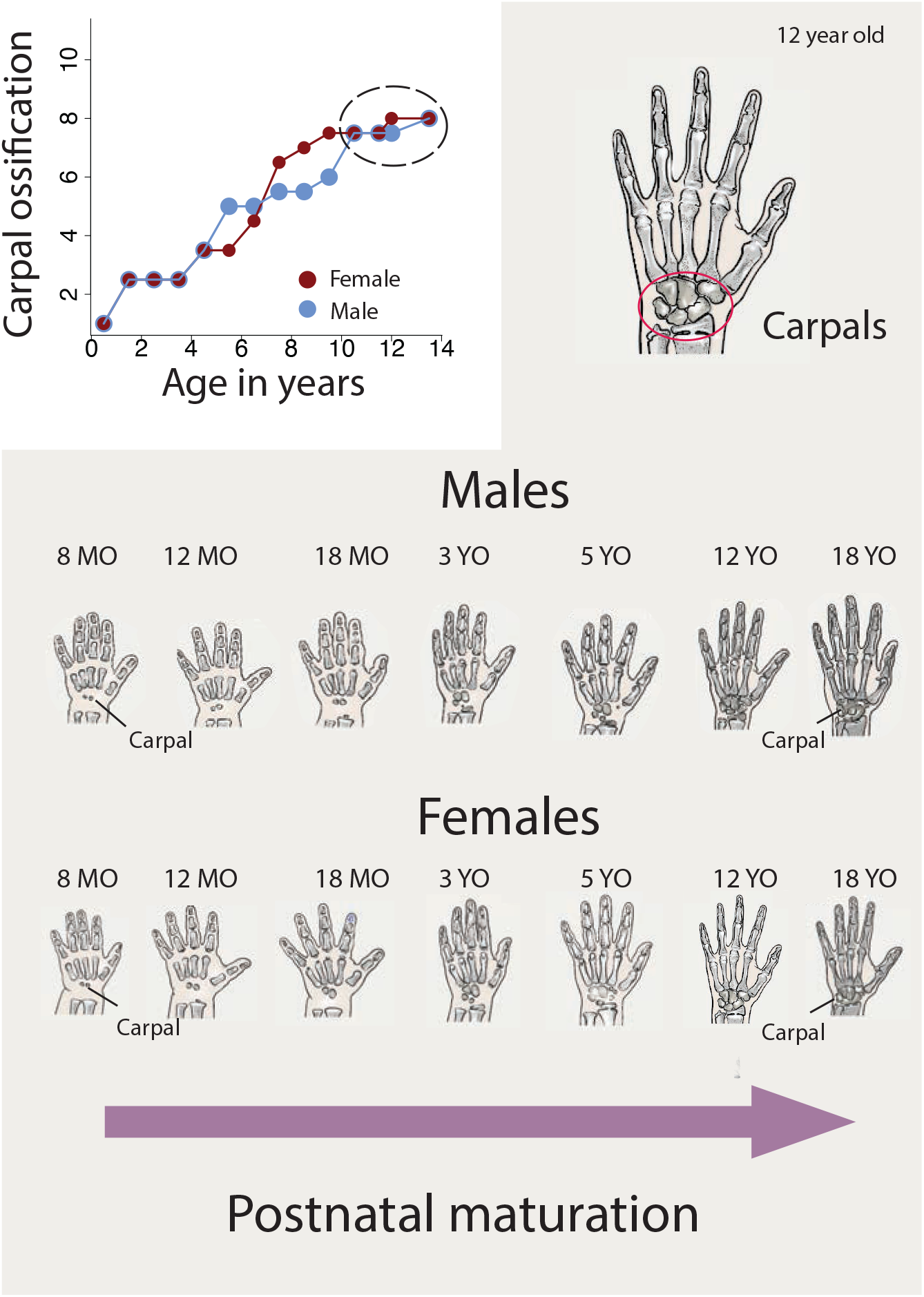
We compared male and female carpal ossification rates in humans. Despite the variation across males and females, carpal ossification is largely complete by 10 to 14 years of age in both sexes. Drawings represent carpal ossification from radiographs of humans at different postnatal ages in male and females in humans and are modified from radiograph atlas (13). There are roughly 2 ossified carpals near birth, and this number steadily but slowly grows postnatally up to around ~12 years of age in humans in males as in females. Abbreviations: MO: months; YO: years. These qualitative observations align with quantitative comparisons (Fig. 7).

**Fig. S17.**
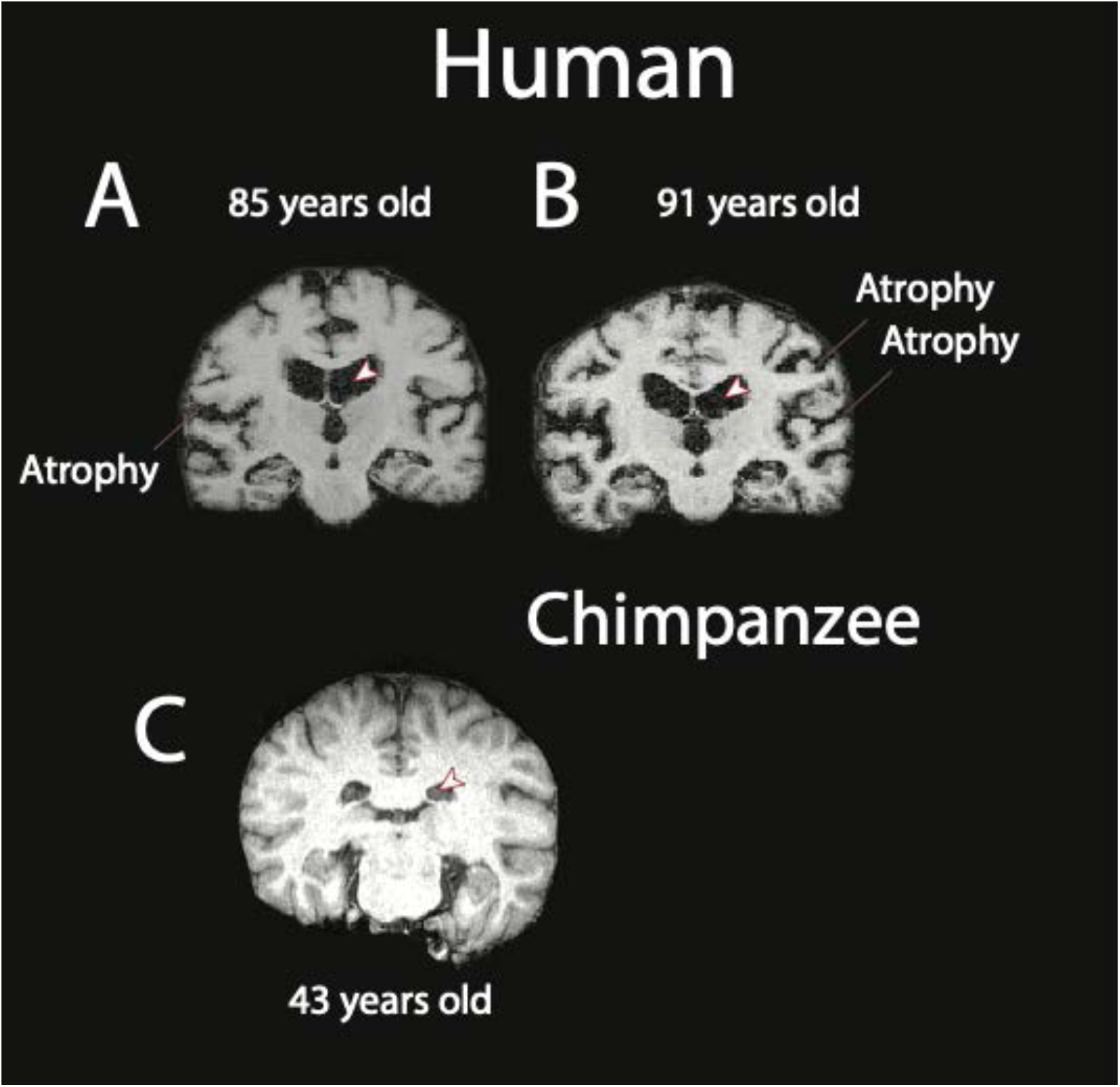
Coronal slices through structural MR scans of aged human and chimpanzee brains show that human brains of 85 and 91 years of age are atrophied, especially at 91 years of age. This is in contrast with aged chimpanzees that do not possess obvious atrophy as exemplified by a 43-year-old chimpanzee. Ventricles (arrowheads) are expanded in aged humans In contrast, these structural modifications with age are not as evident in the aged chimpanzees. Chimpanzee and human structural MR scans are from the National Chimpanzee Brain Resource and the open access series of imaging studies (OASIS) database, respectively (14).

### Supplementary tables

**Table S1.** Time points used to find corresponding ages across primate species.

